# Neuronal activity regulates glycolysis to sculpt synaptic transmission

**DOI:** 10.64898/2026.11.17.766455

**Authors:** Emma K. Theisen, Philip M. Borden, Thomas R. Clandinin

## Abstract

Neuronal computation imposes rapid, localized changes in energy consumption, yet how energy production is regulated at such fine spatiotemporal scales is incompletely understood. Using two photon imaging of neuronal activity and energy metabolism in *Drosophila*, we show that neurons recruit glycolysis in less than a second, with cell-type and subcellular specificity. This coupling scales with neuronal activity and is regulated by AMP-activated protein kinase (AMPK). Activity-induced glycolysis is tightly correlated with presynaptic sites in cholinergic neurons, and is required for cholinergic but not glutamatergic neurotransmission. In cholinergic neurons, glycolysis supports both energetic demand and acetylcholine biosynthesis, with ATP citrate lyase linking these functions by both activating acetylcholine biosynthesis and disinhibiting glycolytic flux. These findings establish glycolysis as a previously unrecognized regulator of cell-type specific computation.

## Main Text

Neuronal computation comes at tremendous energetic expense: brains are metabolically demanding organs, and must accommodate rapid, spatially structured changes in metabolic cost in response to computational need(*1*). These metabolic costs are met by both oxidative phosphorylation and glycolysis, with the latter being particularly well suited for providing a flexible energy supply because of its comparatively fast kinetics. However, how glycolysis might be coupled to neuronal activity over the fast timescales of computation remains unclear, and whether glycolysis might have cell-type specific synaptic functions is unknown.

Energy metabolism in the brain covaries with cell types and subcellular compartments. For example, higher glucose uptake is seen in inhibitory neurons compared to excitatory neurons(*2*), raising the possibility that energy demand could be coordinated with neurotransmitter identity(*2–4*). Intriguingly, neurotransmitter biosynthesis is metabolically coupled to energy producing pathways. For example, glutamate and glutamine recycling intersects directly with glucose metabolism(*5*, *6*), while acetyl-CoA, a product of glycolysis, is required for acetylcholine synthesis(*1*, *7–9*). Recent work further suggests that neuromodulator identity employs distinct metabolic strategies, as dopamine signaling regulates glycogen stores to support presynaptic function in dopaminergic neurons(*10*). At the same time, energy demand is highly compartmentalized within individual neurons. Presynaptic sites are densely populated with synaptic vesicles, which impose substantial ATP demands even at rest(*11*) and still greater demands during activity(*12–15*). Postsynaptic compartments have distinct energy burdens, particularly due to ATP-dependent restoration of ion gradients(*16*, *17*). One way neurons appear to meet these spatially distributed demands is through precise mitochondrial positioning and trafficking(*18–29*). Glycolysis, too, appears to be spatially organized, as glycolytic enzymes cluster at presynaptic sites in response to hypoxia(*30*, *31*), and in vitro studies have described tethering of glycolytic enzymes to synaptic vesicles and mitochondria(*32–38*). More recently, direct imaging of glycolysis in C. elegans neurons revealed that glycolysis is differentially recruited by cell type(*39*). Taken together, these observations raise the possibility that neurons spatially tune energy production to the localized demands imposed by cellular identity and subcellular function. Because energy production is required for neuronal signaling, this metabolic tuning could in turn constrain neuronal function *in vivo*. Yet how glycolysis is spatiotemporally regulated during physiological activity remains unknown.

In this study, we used *in vivo* imaging to capture changes in glycolysis and neuronal activity in the *Drosophila* visual system. We find that neuronal activity is jointly sustained by glycolysis and oxidative phosphorylation, such that either mechanism can compensate for the loss of the other. Using the glycolytic sensor HYlight, we find that neuronal activity rapidly recruits glycolysis in less than a second, a response that varies across neurons and subcellular compartments. In a targeted genetic screen, we identified the AMPK complex as a critical regulator of the coupling between neural activity and glycolysis at short timescales. Correlating glycolytic dynamics with synaptic features using the connectome revealed a strong positive association with presynaptic sites, specifically in cholinergic neurons. Consistent with this organization, we demonstrate that disrupting glycolysis selectively impairs release of acetylcholine, but not glutamate. We find that glycolysis is directly linked to acetylcholine biosynthesis through the conversion of glucose-derived citrate into acetyl-CoA via ATP citrate lyase. As a result, synthesis of acetylcholine promotes glycolytic flux by relieving citrate-mediated inhibition of glycolysis, thereby allowing local tuning of energy production to meet synaptic demand. Thus, we discover that glycolysis is (i) scaled to neuronal activity (ii) flexibly recruited at the level of cell types and subcellular compartments and (iii) specialized not just to meet local energy demands, but also for its biosynthetic roles to satisfy the functional requirements of cholinergic neurons.

### Neuronal activity tunes glycolysis and depends on complementary energy production pathways

Neuronal activity imposes substantial energy demands, suggesting that energy supply must be continuously replenished to sustain neuronal function (Fig. 1A). We therefore first examined whether neuronal activity *in vivo* requires both of the primary forms of cellular energy production, namely glycolysis and oxidative phosphorylation (oxphos; Fig. 1B). To do so, we measured stimulus evoked changes in neuronal and metabolic activity in the *Drosophila* visual system (Fig. 1C). To do this, we leveraged cell-type specific expression of activity and metabolic sensors, and measured responses to flashes of light. Visual signals are transmitted from photoreceptors through the lamina and into the medulla, where distinct neuron types respond selectively to either increases in light intensity, contrast increments, or decreases in light intensity, contrast decrements(*40*). We focused initially on Tm1, a third order interneuron that depolarizes in response to contrast decrements, and measured the responses of individual neurons in a synaptic output compartment denoted M9 (Fig. 1C).

**Fig. 1.**
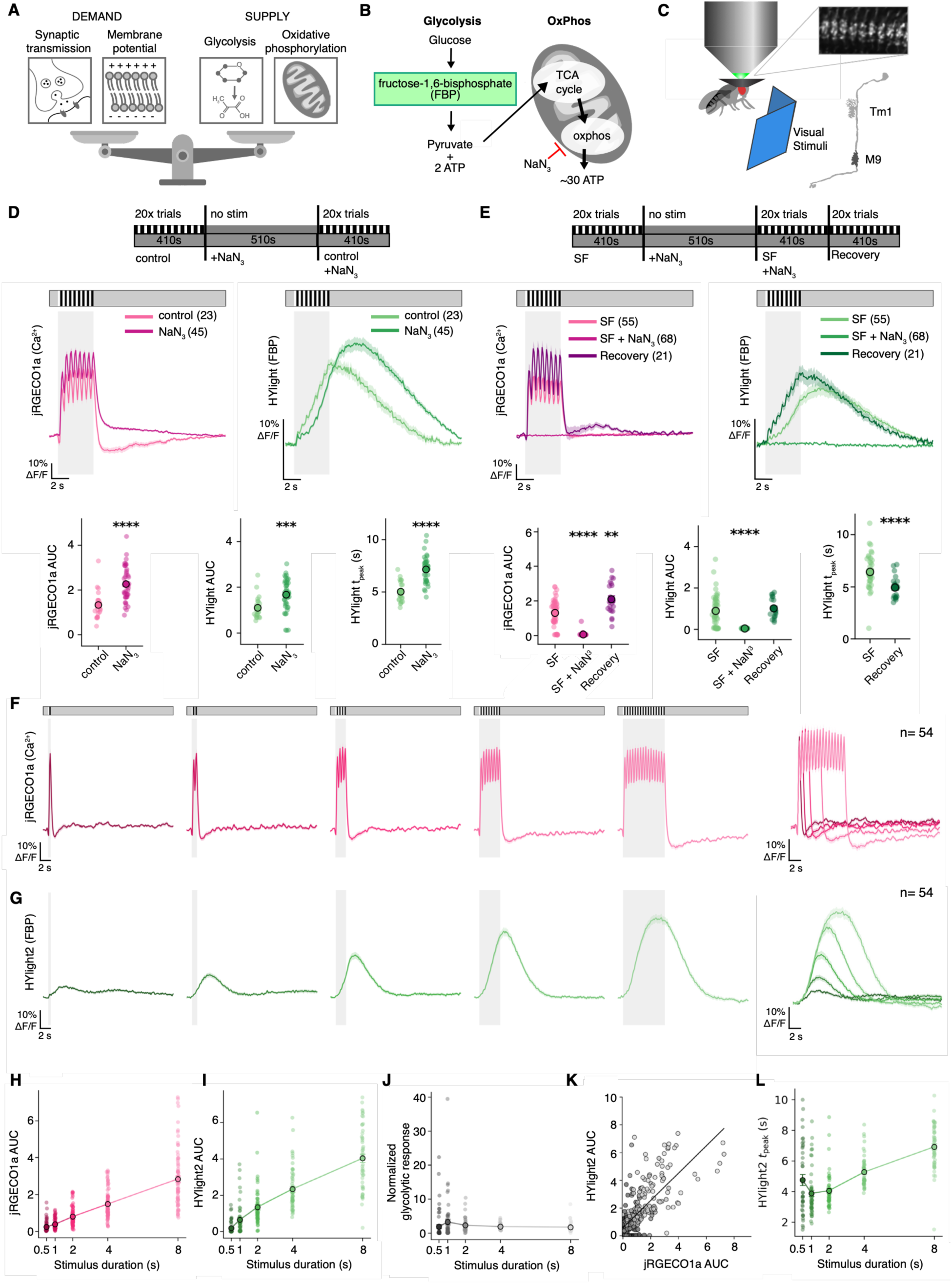
Energy production sustains neuronal activity and glycolysis is scaled to the level of activity demand. (**A**) Schematic illustrating the balance between cellular processes that increase energy demand and those that produce energy in neurons (**B)** Diagram of the two major pathways of energy production in cells: glycolysis and oxphos. Glycolysis breaks down glucose into pyruvate and yields two ATP molecules per reaction cycle. Fructose-1,6-bisphosphate (FBP) is produced during the committed step of glycolysis, which is sensed by the HYlight sensor. Oxidative phosphorylation (OxPhos) takes place in mitochondria and produces ∼30 ATP molecules per reaction cycle. Sodium azide (NaN_3_) is a potent OxPhos inhibitor. (**C**) Schematic illustration of our 2-photon imaging experiment. The fly views visual stimuli presented on a projector screen. Representative M9 terminals from Tm1 are shown in the top right corner, with a schematic of Tm1 in the bottom right. (**D**) Top: Experiment design. Visual stimuli are first presented in control saline followed by NaN_3_ addition over 510 seconds, followed by a second cycle of visual stimulus presentation. Middle: Mean jRGECO1a responses under control and NaN_3_ conditions. Mean HYlight (FBP) recorded simultaneously. Bottom: Area under the curve (AUC) of jRGECO1a and HYlight responses. Each dot represents one neuron. Time-to-peak of the HYlight response. (**E**) Top: Experiment design conducted in sugar free (SF) saline followed by NaN3 addition and a final imaging series during which visual stimuli are presented while recovery saline containing glucose, trehalose, and sucrose is perfused onto the brain. Middle: Mean jRGECO1a responses in Tm1 in SF, SF + NaN_3_, and recovery saline conditions. Calcium responses remain intact after glucose deprivation in sugar-free saline alone (SF), but SF + NaN_3_ eliminates all visual responses. Recovery saline restores jRGECO1a. Mean HYlight responses collected simultaneously. Bottom: jRGECO1a AUC, HYlight AUC and time-to-peak of the HYlight response. N samples (neurons) listed in parentheses. (**F-G**) Mean jRGECO1a and HYlight2 responses to different stimulus durations (**H**) jRGECO1a AUC by stimulus duration. Each dot represents a neuron, with the dark outlined circle representing the mean AUC. (**I**) HYlight2 AUC by stimulus (**J**) Ratio of the HYlight2 response AUC to the jRGECO1a response AUC across stimulus durations (**K**) Each neuron plotted as a function of jRGECO1a AUC (x-axis) and HYlight2 AUC (y-axis). (**L**) Time-to-peak of the HYlight2 response for each stimulus duration. Differences across imaging conditions were assessed using Welch’s one-way ANOVA for groups of two and Welch’s one-way ANOVA for. Experimental genotypes were compared with the pooled control (control + control-RNAi) using two-sided Welch’s t-tests with Holm correction for multiple comparisons. * p <0.05, ** p <0.01, *** p <0.001, **** p<0.0001.

To probe the contributions of glycolysis and oxphos to neuronal activity, we selectively inhibited glycolysis and oxidative phosphorylation (oxphos). To block oxphos, we used the inhibitor sodium azide (NaN_3_) (Fig. 1B, D). To inhibit glycolysis, we perfused the brain with a saline solution lacking glucose, trehalose and sucrose (sugar free; SF; Fig. 1E). Lastly, we performed both manipulations simultaneously(*41*, *42*) (Fig. 1E). Under control conditions, Tm1 neurons expressing the red-wavelength calcium sensor jRGECO1a responded to alternating light and dark flashes with stimulus-locked calcium transients that could be stably maintained over 20 trials (400 seconds)(*43*) (Fig. 1D; fig. S1A-C). Sodium azide caused an immediate increase in intracellular calcium concentrations, consistent with the disruption of neuronal ion gradients during energetic failure, such as during ischemia(*42*, *44*, *45*)(fig. S1A). Intracellular calcium concentrations returned to baseline over 1-2 minutes, and after this recovery period, responses of Tm1 to visual stimuli were enhanced and remained stable for at least 20 trials (Fig. 1D; fig. S1B). These results indicate that while neurons depend on oxphos as an energy source to maintain baseline ion gradients, they can adapt to oxphos inhibition by using an alternate pathway to sustain visual responses. We reasoned that this compensation might reflect upregulation of glycolysis. To test this, we first examined visually evoked responses in the absence of extracellular sugars (sugar free; SF), and observed intact calcium responses (Fig. 1E; fig. S1D-F). However, combining sugar depletion with oxphos inhibition (SF + NaN_3_) abolished visually evoked responses in Tm1 (Fig. 1E; fig. S1E). Strikingly, this effect was reversible, as resupplying extracellular sugars recovered calcium responses that were comparable to controls (Figure 1E; fig S1E). These results indicate that glycolysis and oxphos are redundantly required to sustain visual responses over the timescales probed here.

To assess whether neurons recruit glycolysis to rapidly supply energy during physiological activity, we generated flies expressing the genetically encoded glycolytic sensor HYlight(*46*)(see Methods). HYlight increases in fluorescence when bound to fructose-1,6-bisphosphate (FBP), the metabolite produced during the committed step of glycolysis via the enzyme phosphofructokinase (Pfk). Consistent with prior studies showing increases in FBP in *C. elegans* neurons and in cultured hippocampal neurons over tens of seconds(*39*, *47*, *48*), we found that visually evoked activity also drove increases in FBP, but with surprisingly rapid kinetics. Glycolytic responses began within a few hundred milliseconds after visual stimulus onset and peaked around stimulus offset (Fig. 1D). Thus, glycolysis is engaged on timescales relevant to sub-second neuronal signaling.

We next examined how perturbing energy production impacted glycolytic recruitment in Tm1. Inhibition of oxphos elevated and prolonged the HYlight response paralleling the increased jRGECO1a response, suggesting that glycolysis increases as a function of heightened neuronal activity and energy demand, potentially compensating for a disruption in oxphos (Fig. 1D; fig. S1C). In sugar free saline, HYlight responses were reduced but not abolished, indicating that removal of external sugars does not eliminate activity-associated glycolytic responses (Fig. 1E). During extracellular sugar deprivation, glycolysis may instead rely on mobilization of glucose from intracellular glycogen(*10*, *47*, *49*) or intercellular transfer of substrate from glia (*50*). In contrast, sodium azide application in sugar free saline, abolished calcium and HYlight responses (Fig. 1E). Thus, neurons incapable of responding to visual stimuli with calcium transients were also no longer glycolytically responsive. Strikingly, reintroducing control saline (“recovery”) restored both neuronal activity and FBP responses, indicating that inhibiting energy production temporarily suppressed neuronal activity rather than causing irreversible cell death (Fig. 1E; fig. S1E-F). These data suggest that neurons respond to activity-induced energy demand by rapidly mobilizing glycolysis.

We hypothesized that glycolysis may be actively scaled to the level of neuronal activity. To test this, we used an improved variant of the glycolytic sensor, HYlight2(*51*), with increased dynamic range and sensitivity, and measured jRGECO1a and HYlight2 responses to varying stimulus durations (Fig. 1F-L). Calcium responses scaled reliably with stimulus duration (Fig. 1F). In parallel, glycolytic responses also scaled with stimulus duration, with brief stimuli evoking modest and transient FBP increases, while longer stimuli produced larger and more sustained FBP elevations (Fig. 1G). Quantification of response magnitude as area under the curve (AUC) revealed calcium responses that tracked stimulus duration (Fig. 1H) mirrored by increased HYlight2 responses (Fig. 1I). The same set of stimuli measured with the original HYlight sensor produced similar results, albeit with more modest responses, particularly during the shortest stimuli (fig. S1A-B). We also examined how HYlight2 responses scaled with peak neuronal activity versus total neuronal activity over time. To do this, we used a single light or dark flash of 300ms duration, across a range of contrasts. Tm1 neurons are OFF-selective, meaning that they respond strongly to the onset of contrast decrements and to the offset of contrast increments (fig. S2A-D). While these two types of responses had comparable peak calcium signals, they differed substantially in their duration with offset responses lasting substantially longer than onset responses (fig. S2A-D). Quantifying this pattern revealed that while peak calcium signals poorly predicted the HYlight2 response, the integrated calcium signal was a better predictor (fig. S2A-G). This suggests that the glycolytic response is scaled not by the peak calcium signal, but by its integral.

Finally, to control for signal specificity, we took advantage of the fact that both HYlight and HYlight2 enable ratiometric measurements that can normalize the FBP-bound signal (using two photon excitation at 920nm) with the FBP-unbound signal (excited at 800nm). Because our imaging configuration did not allow simultaneous excitation at both wavelengths, we instead sequentially imaged the same neurons at 920nm and 800nm across all stimulus durations. The resulting 920nm/800nm response curves closely tracked the 920nm FBP-bound signal, indicating that the single-wavelength HYlight response faithfully captured activity-dependent changes in FBP (fig. S3H-Q).

Intriguingly, the time-to-peak of the HYlight2 signal did not scale linearly with stimulus duration, as increased stimulus durations evoked only modest shifts (Fig. 1L; fig. S2H; fig. S2G). This suggests that the timing of the glycolytic response may be governed by a process initiated at the onset of neuronal activity rather than simply by the duration of the stimulus itself. Given that the calcium response and the peak glycolytic response are offset in time by seconds and the timing of the peak glycolytic response is nearly constant across stimuli, these observations are consistent with the notion that neuronal activity is coupled to glycolysis via an intracellular signaling cascade.

### Activity-induced glycolysis is specialized across subcellular compartments and neuron types

Glycolytic enzymes are found in synaptosomes and can localize to synapses during hypoxic stress, raising the possibility that glycolysis may be spatially regulated within neurons (*30*, *39*). We therefore investigated whether activity-driven glycolysis was differentially modulated across compartments and cell types. The *Drosophila* visual system comprises a diverse array of molecularly and morphologically distinct neuron types, each with multiple subcellular compartments. Using cell-type specific drivers, we expressed jRGECO1a and HYlight in eight different cell types, including two second-order interneurons, the lamina neurons L1 and L2, alongside six third-order interneurons in the medulla, namely Tm1, Tm2, Tm3, Tm9, Mi1 and Mi9. Using the 4-second flicker stimulus (Fig. 1), we then measured visually evoked responses across morphologically distinct arbors in each cell type (Fig. 2; fig. S4).

**Fig. 2:**
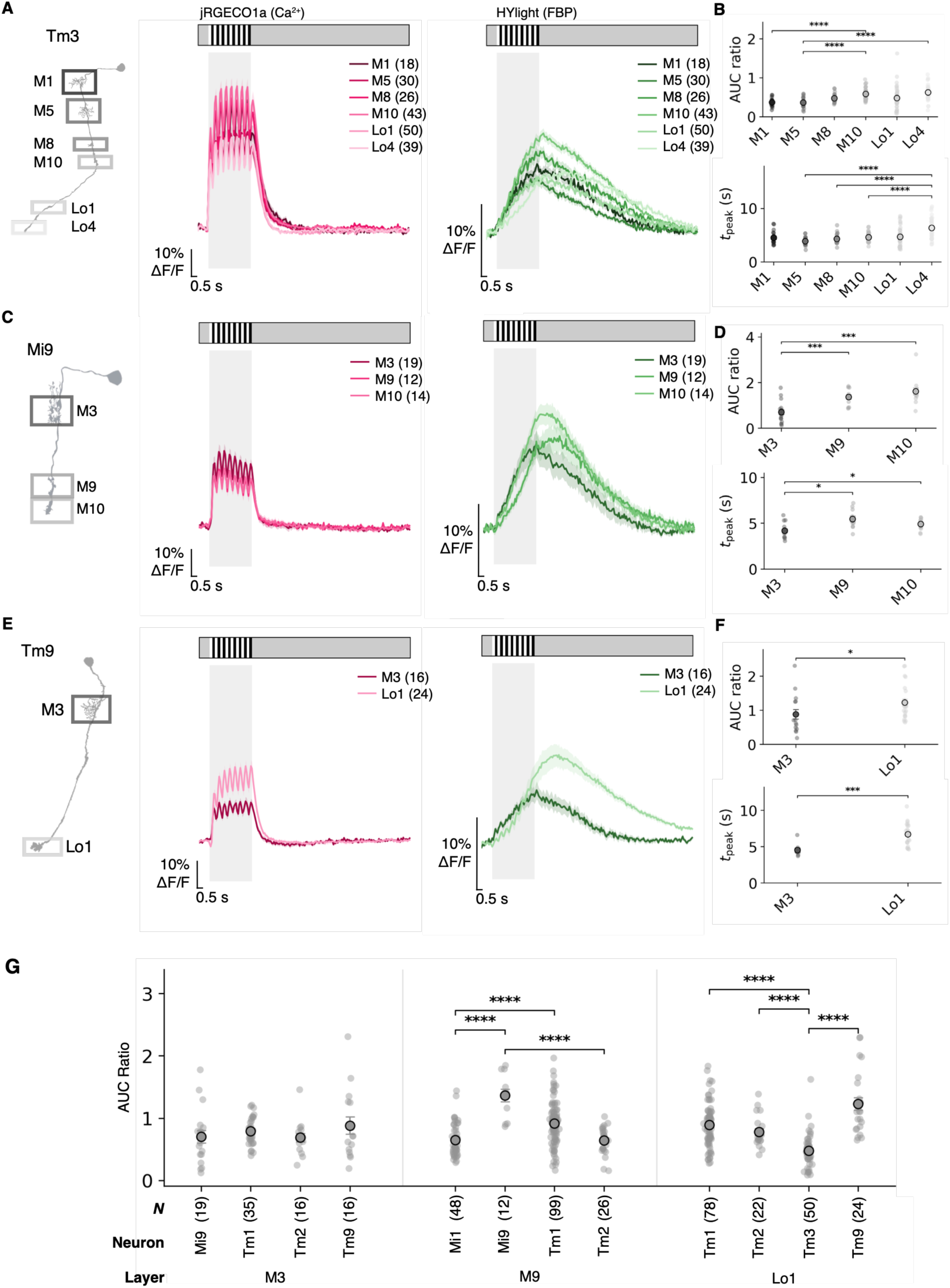
Activity-induced glycolysis varies across subcellular compartments and neuron types. (**A-F**) Subcellular imaging of activity-dependent glycolysis across three medulla neuron types: Tm3, Mi9, and Tm9. Number of neurons per compartment measured listed in brackets in the legend. (**A, C, E**) Left: Illustration of Tm3, Mi9 and Tm9 neurons with separate synaptic compartments outlined by boxes. Middle: Mean jRGECO1a (Ca2+) responses for each compartment of the neuron overlaid, shaded by compartment and mean HYlight (FBP) responses for each compartment and neuron type, shaded by compartment (**B, D, F**) Top: AUC ratio (HYlight AUC/ jRGECO1a AUC) separated by compartment. Bottom: time-to-peak of the HYlight response by compartment. Overall differences across compartments were assessed using Welch’s one-way ANOVA. Experimental genotypes were compared with the pooled control (control + control-RNAi) using two-sided Welch’s t-tests with Holm correction for multiple comparisons. (**G**) AUC ratio (HYlight AUC/ jRGECO1a AUC) across three different anatomical layers that contain synaptic arborizations from at least four neuron types. Kruskal-Wallis test was used followed by Mann-Whitney U tests with Holm-Bonferroni correction across all comparisons (α = 0.05). * p <0.05, ** p <0.01, *** p <0.001, **** p<0.0001.

Across all cell types imaged, we observed increases in glycolytic activity in response to neuronal activity. To quantify the relative strengths of the glycolytic and calcium responses we computed the ratio of HYlight to jRGECO1a of the areas under the curve (AUC ratio), providing a normalized glycolytic response. Intriguingly, in three cell types (Tm1, Tm2, and Mi1) we observed no differences in glycolytic regulation across compartments, while the remaining neurons displayed subcellularly distinct responses. In Tm3 neurons, while the flicker stimulus evoked similar jRGECO1a responses across six compartments (Fig. 2A), HYlight responses in each compartment varied in their time to peak and had distinct amplitudes (Fig. 2B). A qualitatively similar pattern was observed in another cell type, Mi9 (Fig. 2C-D). Using the normalized glycolytic response metric, Tm3 had a relatively small glycolytic response ratio while Mi9 was nearly three-fold larger for their M10-layer compartments (Tm3: 0.58 and Mi9: 1.62). Finally, Tm9 neurons displayed distinct jRGECO1a and HYlight responses across two compartments, with differing time-to-peak values and distinct amplitudes as well as a glycolytic response ratio of comparable magnitude to Tm1 (Fig. 2E-F, fig. S4C).

We next explored whether this glycolytic variation could be accounted for by spatially localized cell-extrinsic factors, such as proximity to trachea and glia, critical sources of energy metabolites. If this were the case, we would expect that the AUC ratio might vary as a function of the anatomical location of each compartment. Three layers, M3, M9 and Lo1, each contain the compartments of four different cell types; if cell-extrinsic factors dominated glycolytic responses, we would predict that responses within a layer, across cell types, would be comparable. However, while the AUC ratio was relatively constant in the M3 layer, M9 and Lo1 displayed substantial variation across neurons, consistent with cell-intrinsic differences (Fig. 2G).

Taken together, by profiling glycolytic dynamics within and across neuron types, these results show that glycolysis is differentially and cell-type specifically recruited across subcellular compartments *in vivo*.

### AMPK acutely couples neuronal activity to glycolysis at short timescales

To understand the functional organization of glycolysis in neurons, we first examined the requirements for each enzymatic step. To do this, we used RNAi to cell type specifically knock down key glycolytic enzymes in Tm1 and measured changes in the normalized glycolytic response (Fig. 3A, fig. S5). As expected, the perturbation of critical enzymes upstream of FBP production (upper glycolysis), namely hexokinase (HexA) and phosphofructokinase (Pfk) substantially reduced the normalized glycolytic response (Fig. 3A-B; fig. S5D). Consistent with the overlap in function between oxidative phosphorylation and glycolysis (Fig. 1), cell autonomous inhibition of Pfk also rendered cells sensitive to oxphos inhibition, resulting in a loss of neuronal response in the presence of sodium azide (fig. S6). In addition, knockdown of genes involved in intracellular glucose uptake and mobilization, namely the glucose transporter (Glut1), trehalase (treh), and glycogen phosphorylase (GlyP), also diminished activity induced glycolysis, but to a lesser degree (Fig. 3A; fig. S5A-C). This suggests that intracellular glucose in neurons is derived from multiple sources. As expected, perturbation of core enzymes downstream of FBP production (lower glycolysis), such as glycerinaldehyde-3-phosphate dehydrogenase (Gapdh3), phosphoglycerate kinase (Pgk), and enolase (Eno) had no effect on activity-dependent FBP production (Fig 3A). However, perturbation of two lower glycolytic enzymes, aldolase (Ald) and pyruvate kinase (PyK), altered the normalized glycolytic response measured using HYlight (Fig. 3A; fig. S5E-F). This suggests that there may be feedback inhibition of upper glycolysis in response to perturbing these key enzymatic steps. Taken together these data demonstrate that activity-dependent glycolysis is a cell autonomous process in neurons.

**Fig. 3:**
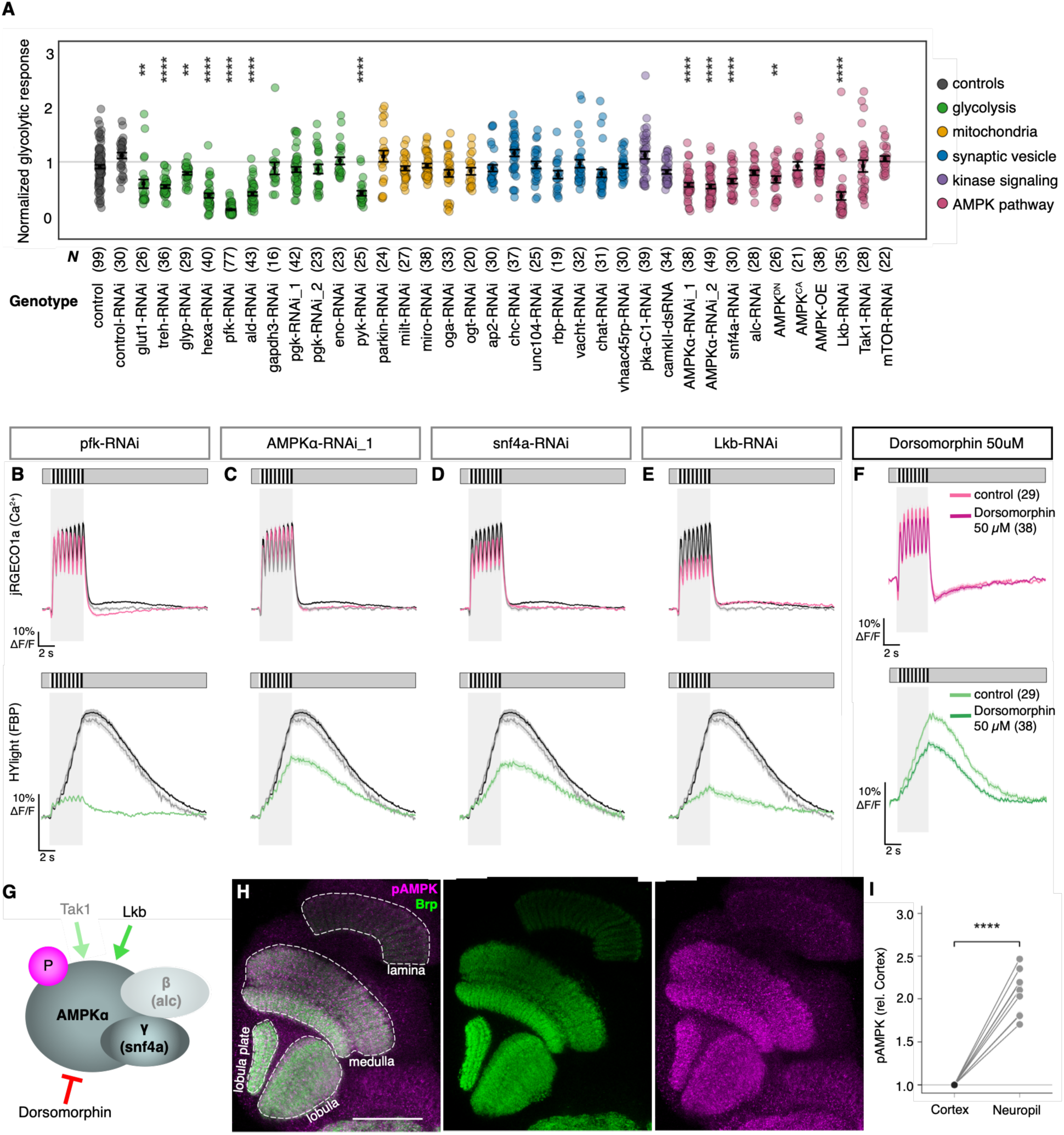
Activity-dependent glycolysis depends on AMPK. (**A**) Normalized glycolytic response plotted for each gene screened. Sample sizes listed represent number of neurons. (**B-E**) Mean jRGECO1a responses in Tm1 expressing the indicated RNAi (magenta), a driver-and sensor-only control (black) and in neurons expressing control-RNAi (gray). (**F**)) Mean HYlight (FBP) responses before (control) and after applying the AMPK inhibitor Dorsomorphin (50uM) (**G**) Schematic illustration of phosphorylated (activated) AMPK (**H**) Immunostaining of phosphorylated AMPK (pAMPK, magenta) co-labeled with synapses (Brp, nc82 antibody, green). Scale bar = 50uM. (**I**) pAMPK immunolabel intensity in synaptic neuropil and surrounding cell-body cortex, with each image normalized to its own cortex value. Lines are individual optic lobes (n = 9). pAMPK is 2-fold higher in neuropil than in cortex (mean +/- SD, paired two-tailed t-test on brain means.) Overall differences across genotypes were assessed using Welch’s one-way ANOVA. Experimental genotypes were compared with the pooled control (control + control-RNAi) using two-sided Welch’s t-tests with Holm correction for multiple comparisons. * p <0.05, ** p <0.01, *** p <0.001, **** p<0.0001.

We next sought to identify a cell-autonomous mechanism that couples neuronal activity to glycolysis. To do this, we performed an RNAi screen in Tm1 neurons and measured the normalized glycolytic response to visual stimuli (Fig. 3A). We first focused on two neuronal functions that have previously been proposed to link activity to glycolysis. First, mitochondria tether glycolytic enzymes and regulate glycolysis in neurons in an activity-dependent manner(*18*, *20*, *32*, *33*). We therefore disrupted mitochondrial localization to synapses by targeting Miro and Milton as well as the adaptor proteins O-GlcNAcase (Oga) and O-GlcNac Transferase (Ogt) that are necessary for tethering. Second, synaptic vesicle recycling is energetically expensive and vesicles tether glycolytic enzymes (*12–15*, *52*). We therefore disrupted vesicle endocytosis targeting the adaptor protein (Ap2), the clathrin heavy chain (Chc), and the vesicular motor protein Unc104. We also targeted vesicle tethering to the active zone by knocking down rim binding protein (Rbp), neurotransmitter biosynthesis (ChAT) and vesicular loading (vAChT) and acidification (VhaAC45RP). However, none of these perturbations had any significant effect (Fig. 3A).

Increased neuronal activity would be expected to consume ATP and increase the levels of ATP breakdown products like AMP. In many cell types, increases in AMP are detected by AMP-activated kinase (AMPK), which in turn activates energy production pathways such as mitochondrial biogenesis(*53*) and silences energy consuming processes(*54*), creating homeostatic feedback loops. However, these known feedback loops are relatively slow, working over minutes to hours, and whether AMPK might work over sub-second timescales to influence glycolysis has not been explored. We therefore knocked down the AMPKalpha and AMPKgamma subunits of AMPK as well as the upstream activating kinase Liver Kinase Beta (Lkb) (Fig. 3A, C-E). Strikingly, all of these perturbations produced substantial reductions in the normalized glycolytic response. Similar results were obtained with expression of a dominant negative AMPKalpha, while expression of a constitutively active form of AMPKalpha had no effect (Fig. 3A). To test whether AMPK is acutely required for coupling neuronal activity to glycolysis, we used pharmacological inhibitors (Fig. 3F; fig. S7). Application of the AMPK inhibitor Dorsomorphin disrupted the normalized glycolytic response, with effect sizes comparable to those seen using cell-type-specific genetic perturbations (fig. 3F; fig. S7A-C) Similarly, BAY-3827, a highly specific AMPK inhibitor(*55*), resulted in a dramatic loss of the normalized glycolytic response, but also impacted neuronal activity, possibly reflecting indirect effects on upstream synaptic inputs (fig. S7D-G). Finally, we note that we did not observe any effect of knocking down either Tak1, the beta subunit of the AMPK complex (alc-RNAi), nor mTOR, one of the prominent regulatory targets of AMPK (Fig. 3A). These negative results could reflect inefficient RNAi or be consistent with the possibility that AMPK has distinct functions at the synapse. Regardless, our results indicate that AMPK is required for the acute coupling of neuronal activity to glycolysis at fast timescales.

Pre- and postsynaptic compartments have distinct energy demands, as presynaptic terminals incur major energy costs associated with synaptic vesicles and postsynaptic terminals use most of their ATP to restore ion gradients(*56*). We therefore sought to identify whether subcellular variation in glycolytic response correlated with structural synaptic features. As an initial effort, we used an antibody that selectively binds the activated form of AMPK, phospho-AMPK, and co-labeled brains with the presynaptic protein Bruchpilot (Brp) (Fig. 3G-I). Consistent with AMPK rapidly regulating glycolysis in axons and dendrites, activated AMPK protein was highly enriched in the neuropil (Fig. 3H-I). This observation is consistent with the observation that AMPK function is required to rapidly couple neuronal activity to glycolytic activity at synapses, and raised the possibility that the glycolytic response might scale with synaptic density.

### Glycolytic responses scale in proportion to presynaptic outputs in cholinergic neurons

To test whether the glycolytic response scaled in association with synaptic features, we took advantage of the connectome, which provides quantitative counts of each presynaptic output and postsynaptic input for every neuron(*57*, *58*). In visual interneurons, subcellular compartments typically contain mixtures of input and output synapses, enabling us to test how the normalized glycolytic response varied as a function of the relative proportion of pre- and postsynaptic sites in each compartment. We mapped pre- and postsynaptic sites in each compartment and normalized each by the total number (see Methods). For example, in Mi1, M9 and M10 are principally presynaptic, while M1 and M5 are primarily postsynaptic (Fig. 4A-C). The total synaptic fraction, representing the sum of pre and postsynaptic sites, also varied by layer, with M1 and M9 having more synapses than M5 and M10 (Fig. 4D). Quantification of synapse distributions for each neuron type in our dataset revealed a broad spectrum of pre- and postsynaptic site distributions across subcellular compartments (fig. S8). Next, we correlated the normalized glycolytic response with these synaptic features. This analysis revealed a positive correlation between presynaptic fraction and the normalized glycolytic response (r = 0.60, p = 0.002), but found no significant correlation with postsynaptic fraction or the total synaptic fraction (Fig. 4E-G). Intriguingly, the sole glutamatergic neurons, Mi9 and L1, were outliers, whose exclusion greatly increased the correlation between presynaptic content and glycolytic response (r = 0.82, p <0.001) (Fig. 4E). These observations raise the possibility that activity-induced glycolysis might play a critical role in presynaptic compartments in cholinergic neurons.

**Fig. 4:**
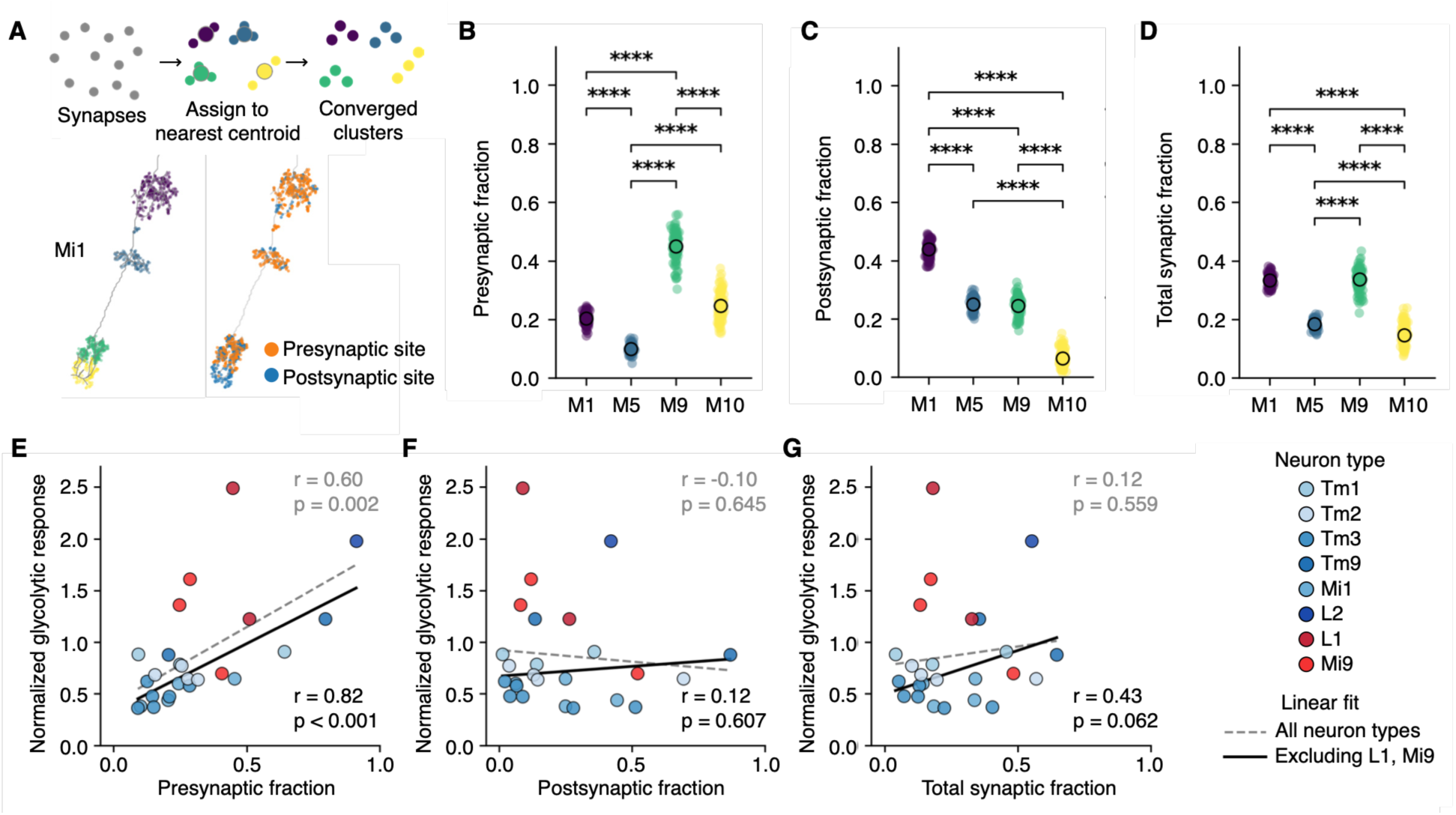
Glycolytic recruitment is associated with presynaptic sites in cholinergic neurons. (**A**)Example Mi1 neuron skeleton with synapses attached from the connectome (neuPrint) and clustered into distinct subcellular compartments using KMeans clustering (top). Bottom left; Synapses are clustered into anatomically distinct compartments, cluster indicated by color. Bottom right: Presynaptic sites are shown in blue, postsynaptic sites are shown in orange. Neuron skeletons and synapse locations were obtained from the optic lobe connectome (dataset optic-lobe:v1.1) (*57*) through neuPrint.(*79*) (**B-G**) Synaptic fractions by compartment for Mi1 (see methods): (**E)** Presynaptic (**F**) postsynaptic and (**G**) total synaptic fraction by compartment. Each dot is one neuron, with n = 100 visually verified neurons. Synapse fractions across compartments were compared within neurons using the Friedman test (Wilcoxon signed-rank test for two compartments), followed by pairwise Wilcoxon signed-rank tests with Holm–Bonferroni correction (α = 0.05). * p <0.05, ** p <0.01, *** p <0.001, **** p<0.0001. (**H-J**) Correlation between the normalized glycolytic response and (**H**) presynaptic fraction, (**I**) postsynaptic fraction, and (**J**) total synaptic fraction of each compartment. Pearson’s correlation coefficient for all cholinergic neurons is positive and significant for presynaptic fraction (black line, r=0.82, p < 0.001) but not other synaptic features at the compartment level. Dashed line and gray r-values indicate Pearson’s correlation calculated for all neurons including the glutamatergic neurons L1 and Mi9 (red).

### Activity-induced glycolysis is required for cholinergic neurotransmission

We next investigated whether the coupling between activity-induced glycolysis and cholinergic presynaptic sites could be due to glycolytic requirements for fueling neurotransmission. Acetylcholine cannot be replenished via reuptake from the synaptic cleft and must therefore be produced *de novo* at the synapse(*59*). Consistent with this, radiolabeling studies have shown that glucose is incorporated into newly synthesized acetylcholine(*60*). We therefore hypothesized that glycolysis may be recruited in an activity-dependent manner to sustain acetylcholine release. To test this hypothesis, we generated transgenic flies expressing a new variant of iAChSnFR that contains a single additional mutation (V188P) and has a higher ΔF/F than iAChSnFR (fig. S9A)(*61*). As the sensor is located extracellularly and increases in fluorescence when bound to acetylcholine, we monitored cholinergic neurotransmitter release in Tm1 neurons expressing iAchSnFR (V188P) (Fig. 5A).

**Fig. 5:**
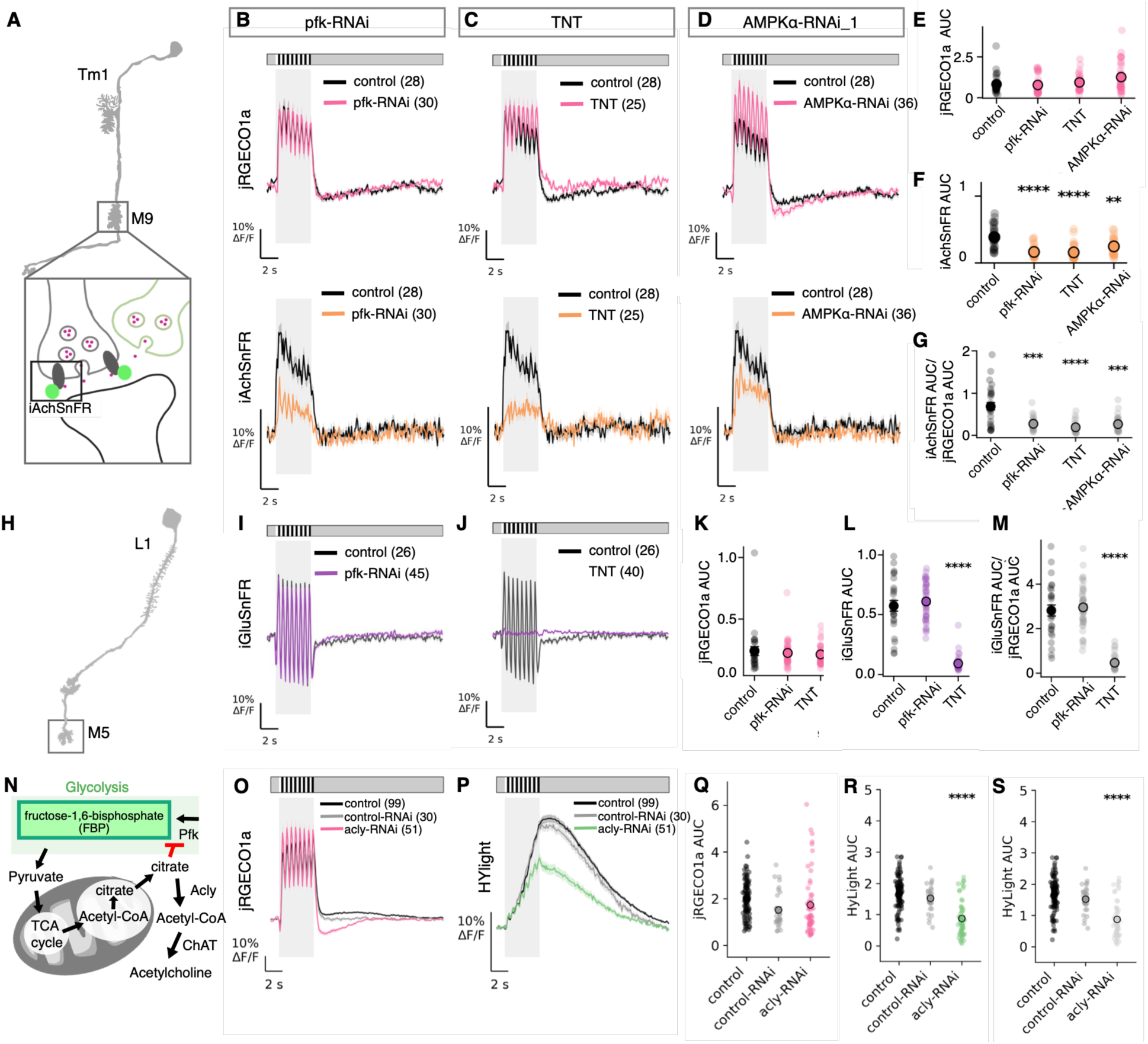
Glycolysis is required for cholinergic neurotransmission. (**A**)Schematic illustration of Tm1. Cutout box shows iAchSnFR (V188P), which fluoresces in response to acetylcholine binding. iAchSnFR (V188P) is agnostic to the source of acetylcholine and can also respond to acetylcholine released by neighboring neurons (represented by the green neuron). (**B-D**) Top: Mean jRGECO1a (Ca2+) responses, with control in black and experimental conditions in magenta. Bottom: Mean iAchSnFR (V188P) (Ach) responses with control in black and experimental conditions in orange. (**E**) jRGECO1a AUC by genotype. (**F**) iAchSnFR (V188P) AUC by genotype. (**G**) The ratio of iAchSnFR (V188P) / jRGECO1a AUC was used to normalize for calcium responses. (**H**) Glutamatergic L1 neurons with the M5 output layer highlighted (**I**) Mean SF-iGluSnFR.A184V measurements in control (black) and pfk-RNAi (purple) conditions. (**J**) Mean SF-iGluSnFR.A184V responses in control (black) and TNT (purple). Quantification of (**K**) jRGECO1a AUC (**L**) SF-iGluSnFR.A184V AUC and (**M**) SF-iGluSnFR.A184V AUC/jRGECO1a AUC. (**N**) Schematic depicting the glycolytic dependence of acetylcholine biosynthesis. Pyruvate produced through glycolysis is used to generate mitochondrial Acetyl-CoA in the TCA cycle, which is converted into citrate before cytosolic import. In the cytosol, citrate inhibits Pfk and is converted into Acetyl-CoA by ATP citrate lyase (Acly), for eventual acetylcholine synthesis by ChAT. (**O-S**) Knocking down Acly results in a dampened HYlight response (**O**) Mean jRGECO1a response (**P**) Mean HYlight responses (**Q**) jRGECO1a AUC (**R**) HYlight AUC and (**S**) normalized glycolytic responses compared between control, control-RNAi and acly-RNAi. Overall differences across genotypes were assessed using Welch’s t-test for two groups and one-way ANOVA for more than two groups followed by two-sided Welch’s t-tests with Holm correction for multiple comparisons. * p <0.05, ** p <0.01, *** p <0.001, **** p<0.0001.

In control Tm1 neurons, iAchSnFR (V188P) responses were elevated during stimulus presentation (Fig. 5B, E-G). To disrupt glycolysis cell-type specifically, we knocked down Pfk in Tm1 neurons (Fig. 4) and observed a significant decrease in acetylcholine release (Fig. 5B, E-G). However, because many neurons in the visual system are cholinergic, the iAchSnFR (V188P) signal could also report acetylcholine release from neighboring neurons (Fig. 5A). We therefore blocked neurotransmitter release from Tm1 via expression of tetanus toxin (TNT), and measured the iAchSnFR (V188P) signal. After TNT expression, iAchSnFR (V188P) responses were dramatically reduced, but not eliminated, thereby revealing contributions from neighboring neurons (Fig. 5C, E-G). Strikingly, this remaining signal was comparable in magnitude to that seen after Pfk knockdown, suggesting that blocking Pfk blocks most, if not all, acetylcholine release from Tm1 neurons. To test whether coupling between neuronal activity and glycolysis is required for cholinergic neurotransmission, we knocked down AMPK and measured acetylcholine release. Notably, knocking down AMPK resulted in a significant reduction of acetylcholine exocytosis, but to a lesser degree than Pfk knockdown, consistent with the effects of these perturbations on the normalized glycolytic response (Fig. 5D-G).

Our connectome analysis indicated that the association between presynaptic content and glycolysis is specific to cholinergic neurons. However, we also observed activity-dependent glycolysis in L1 and Mi9 (Fig. 3, fig. S3), two glutamatergic neuron types, raising the possibility that glycolysis might also be required for synaptic transmission in these cells. To test this, we repeated the knockdown of Pfk and expression of TNT in L1, and measured its neurotransmitter release using SF-iGluSnFR.A184V(*62*). While TNT silenced synaptic release from L1, knockdown of Pfk had no effect (Fig. 5H-M). These results are consistent with the notion that activity-dependent glycolysis is specifically required in cholinergic neurons to replenish acetylcholine pools lost during activity.

Acetylcholine biosynthesis is biosynthetically linked to glycolysis via the TCA cycle, which produces the citrate needed to generate acetyl-CoA in the cytosol via breakdown by ATP citrate lyase (Acly)(*63*). Citrate also provides negative feedback onto glycolysis by inhibiting Pfk(*64*, *65*) (Fig. 5N). We therefore examined whether knockdown of Acly had any effect on glycolysis. Strikingly, Acly knockdown resulted in a significant decrease in the normalized glycolytic response, meaning that citrate flux through Acly is normally used to sustain acetylcholine levels, enhancing the local glycolytic response (Fig. 5O-S). Taken together, our data demonstrate that acetylcholine release requires an activity-dependent glycolytic response, and that it is mechanistically coupled to glycolysis via citrate.

## Discussion

Glycolysis is an evolutionarily ancient metabolic pathway, conserved from anaerobic bacteria to human neurons(*66*). However, how glycolysis might be regulated across different cell types and subcellular compartments, and over fast timescales, remains incompletely understood. Here we show how neurons can rapidly mobilize glycolysis to meet energy demand with exquisite precision. Glycolysis is recruited over sub-second timescales relevant to neuronal computation, scales with the magnitude and duration of neuronal activity (Fig. 1) and is coupled to activity via AMPK (Fig. 3). Recruitment also differs by cell type and subcellular compartment (Fig. 2, fig. S4), and is particularly crucial at cholinergic synapses, where it is necessary for cholinergic neurotransmission (Fig. 4, Fig. 5). These results argue that neurons can optimize energy production strategies to match their computational outputs.

We find that AMPK couples neuronal activity to glycolysis with remarkably rapid kinetics. Previous work has shown that AMPK can regulate glycolysis over longer timescales, by inhibiting glycogen synthesis over tens of minutes(*67*) and by promoting the membrane recruitment of the GLUT3 and GLUT4 glucose transporters over seconds to minutes(*16*, *68*). However, these mechanisms do not readily explain the rapid glycolytic tuning we observe, because *Drosophila* lacks a GLUT4 ortholog and visual system neurons do not express GLUT3(*69*). We instead favor a model where synaptically localized AMPK acts as a control node that senses local, transient increases in AMP. Activity-dependent ATP consumption, particularly associated with synaptic vesicle recycling(*12–15*) and ion gradient restoration(*16*, *17*), can locally increase AMP and ADP relative to ATP and thereby activate AMPK. This would provide a direct feedback mechanism for coupling glycolysis to energy demand(*54*). Our genetic perturbations reveal that nearly all of the canonical members of the AMPK complex regulate activity-induced glycolysis, consistent with a direct activation mechanism caused by increases in ADP and AMP(*70*, *71*). Downstream, AMPK may activate 6-phosphofructo-2-kinase (Pfrx in *Drosophila*), which contains AMPK target sites and produces fructose-2,6-bisphosphate, a potent activator of Pfk(*72*). Rapid modulation of Pfk would provide an attractive mechanism for modulating glycolysis over sub-second timescales, because it would not require new protein synthesis, transporter localization, or large-scale redistribution of metabolic machinery.

Our data also show that neuronal glycolysis is regulated hyperlocally. Prior studies have identified metabolic differences between neuron types(*2*, *10*) and between the soma and neurites in C. elegans sensory neurons(*39*). We build on this work by resolving activity-induced glycolysis within 25 anatomically defined sub-compartments across eight different neuron types (Fig. 2, fig. S4) and find that glycolytic recruitment differed substantially across neurons and subcellular compartments. Whereas connectome studies have mapped mitochondrial density and localization as static structural proxies for metabolic capacity(*24*, *25*), we show that the connectome can also provide a scaffold for mapping metabolic function onto neuronal architectures. In particular, activity-induced glycolysis was associated with presynaptic specializations, consistent with glycolytic enzymes tethering on synaptic vesicles(*34*, *37*, *38*) and clustering at presynaptic sites during hypoxia(*30*, *31*). Presynaptic compartments also incur high energy costs through functions like synaptic vesicle recycling and maintaining synaptic vesicle supply(*11–13*, *56*). Locally enhancing glycolysis at presynaptic locations may therefore represent an economically efficient strategy for optimizing energy production. Intriguingly, presynaptic architecture was most strongly able to predict glycolytic demand in cholinergic neurons. Thus, synapses cannot be assigned uniform energy costs as these costs differ by neurotransmitter identity. Different modes of neurotransmission may therefore recruit distinct metabolic strategies.

Most prior work on neuronal energy metabolism has focused on cultured mammalian neurons, which are predominantly glutamatergic, leaving comparatively little known about metabolic regulation in cholinergic neurons. One unexpected finding from our work was the selective dependence of cholinergic neurotransmission on glycolysis. This dependence reflects a distinction between glycolytic burden and glycolytic sensitivity, a principle previously proposed in the context of presynaptic energetics. Under this principle, processes most sensitive to reduced ATP levels, such as synaptic vesicle endocytosis, are not necessarily those that impose the greatest energy burden(*13*). Activity-induced glycolysis was not uniquely elevated in cholinergic neurons relative to glutamatergic neurons, yet disrupting glycolytic flux selectively impaired cholinergic transmission. Conversely, perturbing acetylcholine synthesis or vesicular loading did not change glycolytic responses (Fig. 3A), arguing that acetylcholine production does not impose a substantial glycolytic burden. Thus, cholinergic transmission depends on glycolytic flux even if acetylcholine production itself accounts for only a small fraction of the glycolytic response.

This tight coupling may arise because acetylcholine, unlike glutamate, must be synthesized de novo, cannot be recycled(*73*, *74*), and its synthesis requires acetyl-CoA derived from glycose through pyruvate and citrate. Citrate is both a precursor for cytosolic acetyl-CoA and an allosteric inhibitor of Pfk^71,72^, and citrate levels transiently decrease during neuronal activity, suggesting a plausible disinhibitory mechanism for enhancing activity-induced glycolysis(*48*, *75*). Indeed, ATP citrate lyase is required to maintain glycolytic responses (Fig. 5), consistent with elevated citrate inhibiting Pfk when its conversion to acetyl-CoA is blocked, and arguing that this conversion supports acetylcholine synthesis in these neurons. Consistent with this biosynthetic link, enhancing glycolysis in cholinergic neurons increased acetylcholine levels in the mouse neuromuscular junction(*9*), and inhibition of ATP citrate lyase decreased glucose incorporation into newly synthesized acetylcholine in rat brains(*60*). Cholinergic neurons may therefore be distinguished not by an unusually large glycolytic burden, but by a particularly tight coupling between glycolytic flux and neurotransmitter production. These results elucidate how glycolysis can multitask to simultaneously fulfill biosynthetic demands and energy demands.

The specialized dependence of cholinergic neurotransmission on aerobic glycolysis may represent a specific vulnerability for cholinergic neurons. In mammals, cholinergic basal forebrain neurons project extensively across the brain and are particularly vulnerable to degeneration in Alzheimer’s Disease (*76*, *77*). Intriguingly, glucose metabolism also declines in Alzheimer’s Disease patients and, more broadly, deficits in energy metabolism have been linked to neurodegenerative disease (*7*, *56*, *78*). Combining these findings with our data and with prior work linking glucose metabolism to acetylcholine synthesis (*9*, *60*), suggests a new vulnerability in these cells, given the obligate relationship we describe between glycolysis and acetylcholine release. At a higher level, our work suggests a general principle of metabolic flexibility as a core feature of neuronal information transmission. We find that metabolic functions of glycolysis and the TCA cycle can support cell-type specific neurotransmission, providing new potential connections between energy metabolism and brain function, with implications for neurodegenerative disease. Exploring how these regulatory mechanisms vary across cell types, circuits, and behavioral contexts is likely to provide new insights into how the brain computes energy efficiently.

## Acknowledgments

We thank Liqun Luo, Lorna L. Jayne, and John P. Vaughen for comments on the manuscript; Manze Zhang and Michelle Y. Pang for fly resources; Yukun Alex Hao for python code to motion correct 2-photon imaging data; Maxwell H. Turner for providing initial guidance for 2-photon microscopy and resulting analysis; Members of the Clandinin Lab and Colleen N. McLaughlin for feedback on the project. TRC is an investigator with the SF Biohub.

## AI use declaration

AI assistance from Claude code (Anthropic) was used to support the generation and debugging of Python code. After using these tools, the authors reviewed and verified the code to be working as intended and take full responsibility for the content of the publication.

## Author contributions

Conceptualization: EKT, TRC

Methodology: EKT, PMB, TRC

Investigation: EKT Visualization: EKT

Formal analysis: EKT

Writing–original draft: EKT

Writing–review and editing: EKT, TRC

Funding acquisition: TRC

Supervision: TRC

## Data, code, and materials availability

All datasets generated in this study are deposited in the Dryad repository. All software and analysis code used in this study can be found on GitHub (see Materials and Methods). All flies generated for this study are available upon request.

## Supplementary Materials

### Materials and Methods

Figs. S1 to S10

### Materials

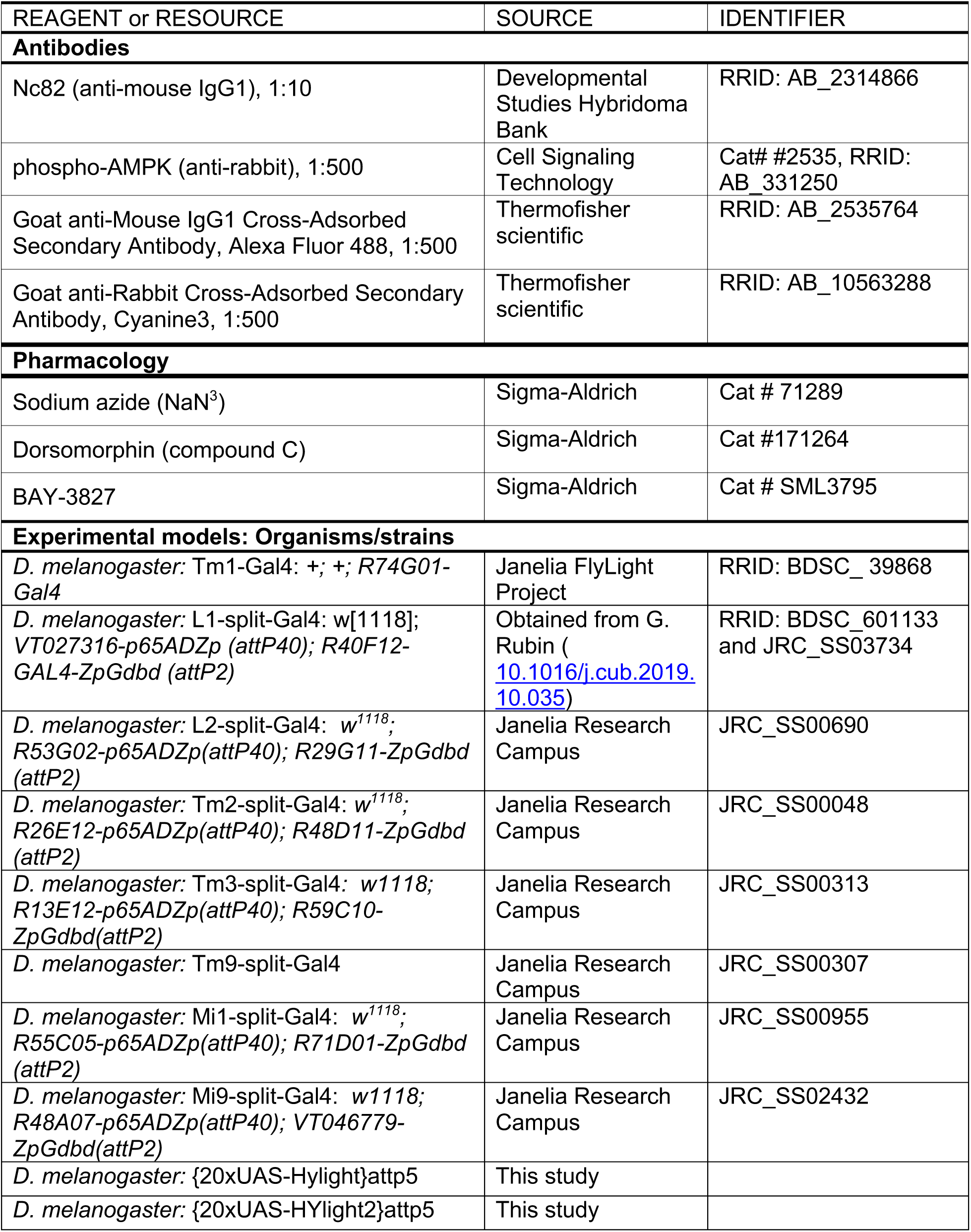
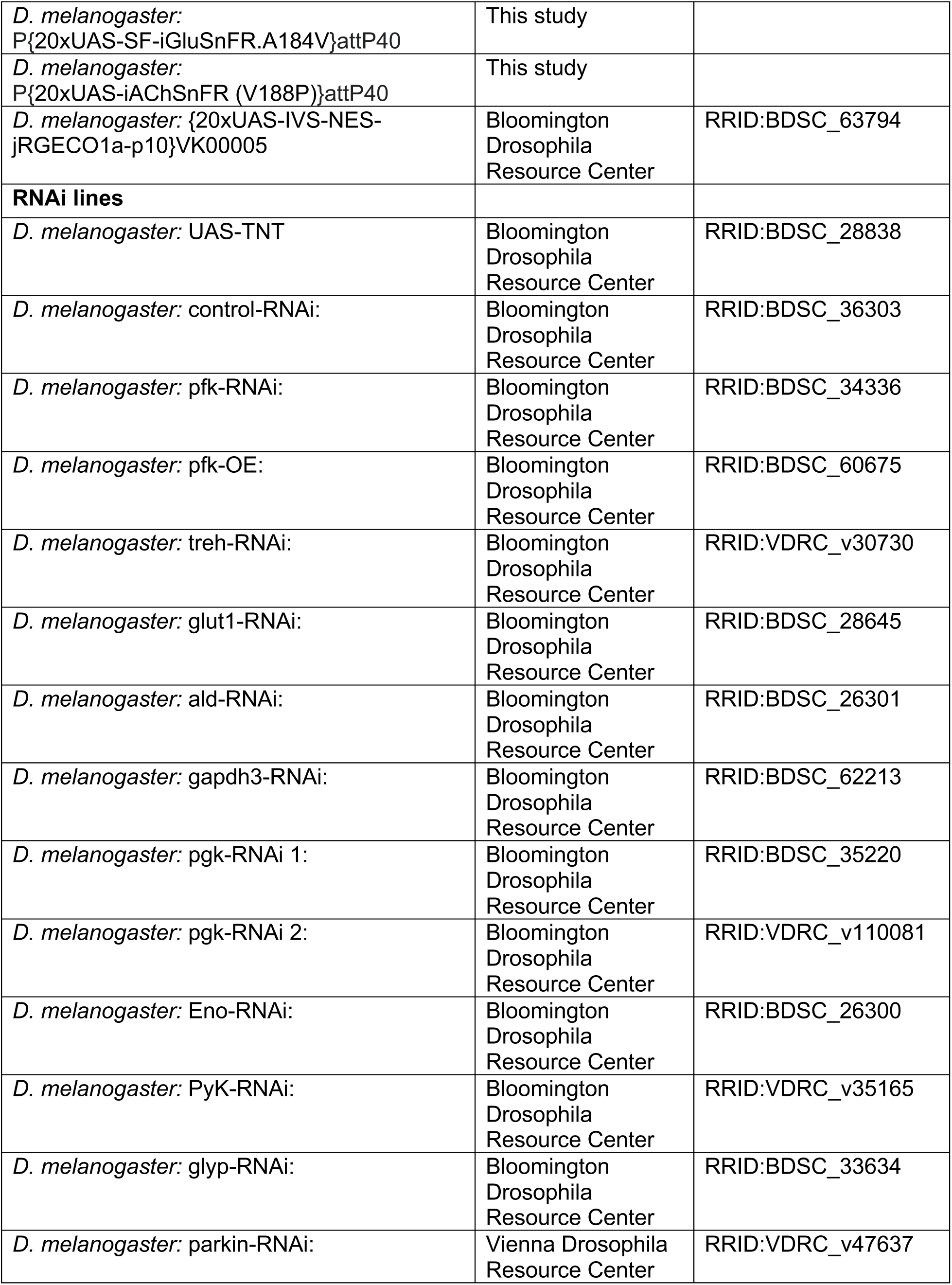
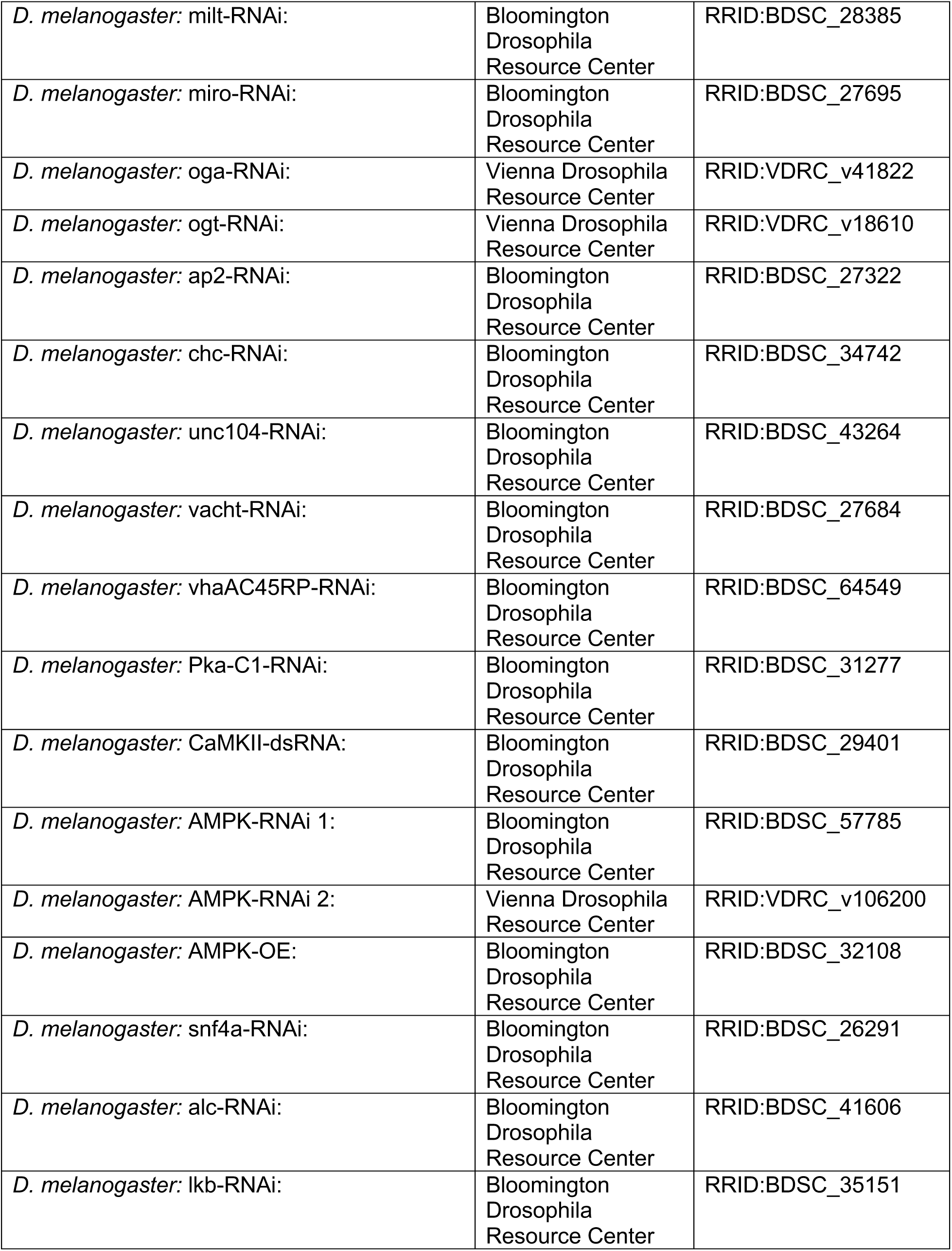
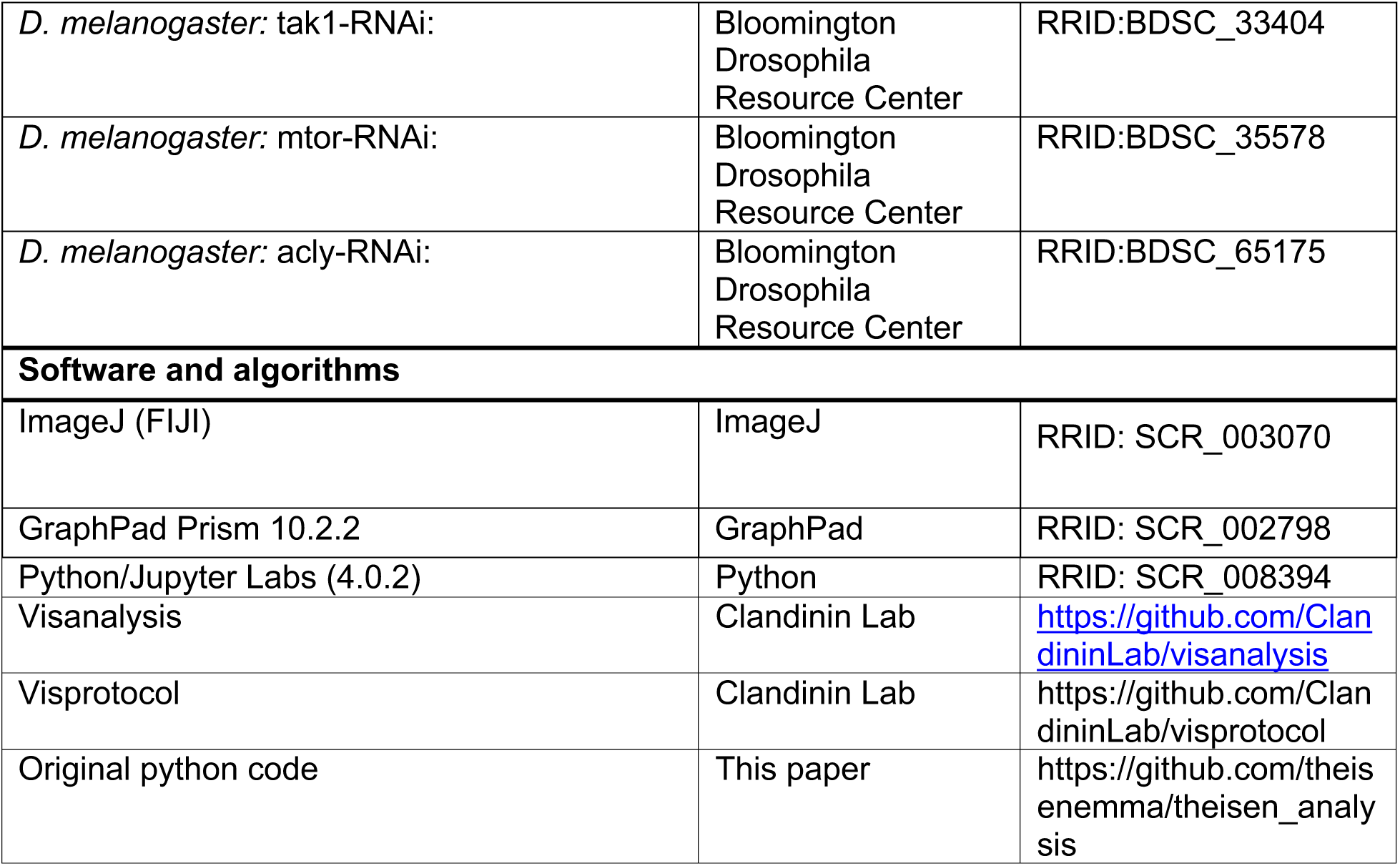

## Methods

### *Drosophila* Maintenance

Flies were grown in vials containing standard molasses-based food and breeding crosses for experiments were kept in 12:12 light/dark (LD) incubators at 25°C with 50%-70% relative humidity. Female flies were imaged between 4-12 days old, with the following exceptions: (1) L1 SF-iGluSnFR.A184V imaging experiments, in which case flies were 1-4 days old to accommodate early adult lethality of TNT expressed in L1 split GAL4 drivers and (2) flies imaged for the RNAi screen were at least 5 days old post-eclosion to facilitate sufficient expression of the RNAi and increase knockdown potency. In all experiments, females were heterozygous for the white gene (or wild-type for the white gene) to avoid behavioral deficits associated with loss of the white transporter in *w-*backgrounds.

### Molecular Genetics

We used FlyBase to find information on stocks, phenotypes, and sequences^91,92^.

The HYlight sequence from Koberstein et al. was codon optimized for *Drosophila*, synthesized (Twist Bioscience), and cloned into the pJFRC7-20X-UAS vector (Addgene #26220). UAS-HYlight plasmids were injected by BestGene Inc and integrated into attp5 (chromosome II) via phiC31 integrase mediated recombination.

Flies expressing 20xUAS-iAChSnFR (V188P) and 20xUAS-SF-iGluSnFR.A184V were generated using the following protocol: the cDNA sequence of each construct was inserted into the pJFRC7-20XUAS vector (Pfeiffer et al., 2010) (Genscript Biotech). Transgenic flies were generated by PhiC31-mediated integration of the construct to produce landing site insertions at attp40 (BestGene): P{20xUAS-SF-iGluSnFR.A184V}attP40, and P{20xUAS-iAChSnFR (V188P)}attP40. Both construct insertions express well.

### Immunohistochemistry

Wild-type isoD1 flies were cold-anesthetized before brain dissection in dissection saline (103mM NaCl, 3mM KCL, 5mM TES, 1mM NaH2PO4, 4mM MgCl) before fixation in 4% paraformaldehyde for 20 minutes. Brains were washed three times in 0.5% PBST (TritonX-100 0.5% in 1X PBS) and placed in blocking buffer (10%NGS) for at least four hours in 96-well plates. Primary antibodies were diluted in blocking buffer and brains were incubated in primary antibody at room temperature for 3 hours followed by overnight incubation at 4°C. Before switching to secondary antibody, brains were washed with 10% PBST three times for 20 minutes. Brains were incubated in secondary antibody at room temperature for 3 hours followed by overnight incubation at 4°C. Finally, secondary antibody solution was removed and brains were washed with PBST three times for 20 minutes each before placement in Vectashield.

The following primary antibodies were used: nc82 (mouse IgG1, 1:10, DSHB, RRID = AB_2314866), phospho-AMPK (Thr172) (rabbit, 1:500, Cell Signaling Technology #2535, RRID = AB_331250). The sequence surrounding Thr172 is fully conserved in *Drosophila* AMPKalpha, in which the equivalent residue is Thr184^93^. The secondary antibodies used were: goat anti-mouse IgG1 AlexaFluor 488 (1:500, Thermofisher, #A21121, RRID = AB_2535764) and goat anti-rabbit IgG Cyanine 3 (1:500, Thermofisher, #A10520, RRID = AB_10563288).

For confocal imaging, brains were placed in Vectashield mounting solution on microscope imaging slides and oriented in dorsal view for optic lobe imaging underneath a coverslip. Optic lobes were next imaged on a Leica SP8 confocal using a 40X Lens (N.A. 1.30) at 1.25x digital zoom with z=2um. Stacks were acquired sequentially so as to prevent bleedthrough signal.

### *In vivo* two-photon imaging

Neurons were imaged following previously described methods^94,95^. We cold anesthetized female flies and tethered them to aluminum shims in fly-shaped holes using UV curing glue. We continuously perfused carbogen-bubbled imaging saline over the fly brain throughout imaging. Imaging saline was kept at 25°C temperature. Unless otherwise noted the imaging saline was prepared with the following recipe: 103 mM NaCl, 3 mM KCl, 5 mM TES, 1 mM NaH2PO4, 4 mM MgCl2, 1.5 mM CaCl2, 10 mM trehalose, 10 mM glucose, 7 mM sucrose, and 26 mM NaHCO3. The pH of the saline equilibrated near 7.3 when bubbled with 95% O2/5% CO2. Sugar free experiments were imaged in imaging solution prepared with the same recipe but omitted the trehalose, glucose and sucrose.

Imaging was performed using a Bruker two-photon resonant-scanning microscope with a 20x, 1.0 NA objective (Leica). To collect calcium responses, jRGECO1a was excited at 1070nm, while all other sensors were excited at 920nm, using ∼15mW post-objective power. Fluorescence signals were collected through a 525/50nm emission filter (Semrock). Before recordings, a search stimulus was used to identify neurons with receptive fields in the center of the projector screen. Only recordings exhibiting clear jRGECO1a responses to the presented visual stimuli were included.

### Pharmacology experiments

For experiments where we inhibited oxphos with NaN_3_, we injected 50mM solution of NaN_3_ into the imaging chamber containing imaging saline for a final concentration of 2mM. Sugar-free saline was the same as the imaging saline, except without trehalose, glucose, or sucrose included. Addition of NaN3 caused an initial decrease in jRGECO1a fluorescence in the terminals, followed by a large increase. To average responses across experiments, traces were aligned to the minimum of this initial decrease (t=0) and plotted as ΔF/F with baseline F0 from the mean trace −10 to −2 seconds relative to the jRGECO1a minimum (t=0) (fig. S1, fig. S6).

For AMPK inhibition experiments, we similarly added Dorsomorphin and BAY-3827 into the imaging dish to final concentrations of 50uM for Dorsomorphin and 35uM for BAY-3827. While Dorsomorphin and BAY-3827 were first solubilized in DMSO as stock solutions, these stock solutions were diluted down in imaging saline for a final concentration of 0.5% DMSO for Dorsomorphin and 0.17% DMSO for BAY-3827 before adding.

### Visual stimulation

Visual stimuli were projected using the blue LED of two DLP LIghtCrafter 4500 projectors onto a fabric screen covering the visual field of the fly. We used a 482/18nm bandpass spectral filter to limit bleed through. We used two open-source software packages developed in the lab (https://github.com/ClandininLab/) – visprotocol to generate and present visual stimuli for the fly and visanalysis to analyze neuron responses to the stimuli presented.

#### Flickering light flash stimulus (Fig. 1, Fig. 2, Fig. 3, Fig. 5)

Light and dark flashes lasting for 250ms per flash were presented during a four second stimulus window, with one second of pre-stimulus time followed by 15 seconds of post-stimulus time. Each trial duration lasted 20 seconds and was presented 20 times. The mean of signal measured in the one second immediately before the first flash was taken as the pre-stimulus period and used as the baseline F in ΔF/F calculations.

#### Duration series (Fig. 1)

The duration of light and dark flashes (each lasting 250ms, and being of full contrast) was randomly interleaved and selected between five stimulus durations: 0.5 seconds, 1 second, 2 seconds, 4 seconds, and 8 seconds. Each stimulus started with a light flash as with the single stimulus series (with 4 second stimulus duration). The total trial duration time was extended to 30 seconds accommodate the 8-second long flash.

#### Duration series for neurotransmitter release imaging (Fig. 5)

To examine neurotransmitter release, we presented four durations: 0.5, 1, 2, and 4 second-long stimulus trials. These were randomly interleaved at 20 second intervals.

#### Short flash stimulus (fig. S2)

Alternating 300ms full-field light and dark flashes ranging across six Michelson contrasts of 0.1, 0.3 and 0.5 were presented in random order with 15-second long inter-stimulus intervals. Each fly was presented with 50 trials of this stimulus and the trial averages across neurons and flies were plotted. Peak ΔF/F was calculated as the maximum rolling average over a 100ms time window during the stimulus period.

### Analysis

Data visualizations and plotting functions were written using Python scripts based on functions available in the visanalysis package, with some plotting functionalities added with the assistance of ChatGPT (OpenAI) and Claude (Anthropic). All generated code was manually verified and tested to ensure reproducibility and faithful derivation from the underlying data.

Regions of interest (ROIs) in each neuron were manually selected in the time-series averaged image in the visanalysis GUI restricted to each synaptic compartment, as represented in Fig. 1C.

### AUC calculation

For all experiments, with the exception of impulse responses with 300ms light and dark flashes (Figure 2, Figure S2), AUC was calculated as the absolute value of area of the region bounded by the horizontal axis and the response using the trapezoidal rule. The vertical margins of the area were defined at x=1 (onset of the stimulus) and the first zero crossing after the stimulus termination was used the rightmost boundary. For the impulse responses, we used the same method, but integrated the first phase of the positive AUC only, because we observed biphasic responses. All AUC measurements were visually verified.

### Time-to-peak

The time-to-peak was calculated after low-pass filtering (butterworth) ΔF/F data to 4Hz. The peak was defined as the maximum value within 15 seconds after stimulus onset, and time-to-peak was recorded as the latency of time in seconds from stimulus onset to the peak value. Only traces whose peak exceeded a fixed amplitude threshold (ΔF/F >= 0.12 for jRGECO1a and >= 0.06 for HYlight) were included so as to prevent noise artefacts.

### Ratiometric normalization of HYlight and HYlight2

For ratiometric normalization of 920nm/800nm acquired wavelengths, background-subtracted fluorescence traces were first independently converted to ΔF/F for 920nm and 800nm excitation using the same pre-stimulus 1-second baseline window. Because 920nm and 800nm excitation trials were acquired sequentially (in random order) and stimulus durations were randomly interleaved, ratiometric traces were computed after grouping trials by stimulus condition. For each ROI and condition, trial-averaged normalized fluorescence traces were computed as 1+ ΔF/F and the 920nm trace was divided by the corresponding 800nm trace per cell level. The resulting ratio was baseline-normalized to generate ΔR/R.

### Quantifying compartmentalized synapses using the connectome

Neuron skeletons, synapse locations and soma locations were obtained from the optic-lobe:v1.1 neuPrint connectome dataset (*64,86*). For each cell type, except for L1 and L2, we analyzed 100 neurons that had a soma location recorded and were not cropped at the volume edge. As only neurons with soma annotations were included, this resulted in fewer than 100 averaged neurons for L1 (n = 90) and L2 (n = 78). For each neuron, healed neurons were fetched and these were visually verified to ensure all neurons included were fully reconstructed and had the expected stereotyped architecture for the given neuron type. Annotated pre- and postynaptic sites were obtained using fetch_synapses and assigned to its nearest skeleton node.

To measure synapse positions faithfully along the neuron skeleton for the quantification of synapses in compartments, each skeleton was represented as a graph, with edges weighted by segment length. Distances along the axon were computed using Dijkstra’s shortest path algorithm. The soma was assigned to its nearest skeleton node and the distance of each synapse from the soma along the neuron skeleton was calculated. The main axis of the neuron was defined as the longest path along the neurite between two synapse-bearing nodes starting at the synapse closest to the soma. Each synapse was assigned a position on its axis scaled from 0 (start, closest to soma) to 1 (end, furthest from soma).

For most neurons, synapses were assigned to compartments by KMeans clustering of their x/y/z coordinates into N clusters. N was specified based on the known anatomy of medulla neurons aligning with the morphology of our imaging data ROIs. Cluster labels were then reordered by the median projected trajectory coordinate of synapses within each cluster, producing a consistent proximal-to-distal ordering across neurons. Cluster assignments were manually inspected for every neuron to ensure correspondence with anatomical optic lobe layers.

For Tm3, KMeans clustering did not faithfully separate synapses by anatomical layer. Tm3 synapses were therefore assigned to compartments using projected-gap clustering along the normalized trajectory coordinate. Synapses were ordered by their trajectory position, and compartment boundaries were defined by large gaps in this ordered distribution. Candidate gaps were required to exceed 1.5 times the median inter-synapse gap, and clusters were required to contain at least 15 synapses. This was set to return six ordered clusters per Tm3 neuron, and cluster labels were reordered from proximal to distal based on mean trajectory coordinate.

Ordered clusters were mapped to named anatomical compartments according to their position within the optic lobe layers. For each neuron and compartment, presynaptic and postsynaptic sites were counted separately by summing synapse annotations classified as pre or post, respectively. Total synapse number was calculated as the sum of presynaptic and postsynaptic sites within each compartment. Compartment-level synapse counts were then used for downstream comparisons with compartment-specific functional imaging measurements.

### Correlating subcellular synaptic features with subcellular glycolytic response

To enable correlation analysis between synaptic features and the normalized glycolytic response across the 25 synaptic compartments we imaged, we used the mean normalized glycolytic response and the mean of each synaptic feature obtained from the connectome and calculated Pearson’s r between these features. The correlation scatterplots were plotted with least squares linear fit lines. Synapse clusters were assigned to compartments separately for each neuron based on their distance from the soma (for example in Tm1, cluster 0 was M2 and cluster 3 was Lo1). To normalize for synaptic content within each neuron, the presynaptic, postsynaptic, and total synapses in each layer were divided by that neuron’s total pre-, postsynaptic and total synapses across all layers for each feature. These fractions were then averaged for the neurons of each type.

### Quantification of pAMPK enrichment in synaptic neuropil

The ratio of mean pAMPK intensity in the neuropil over the mean pAMPK intensity in the cell-body cortex was used as a measurement of pAMPK enrichment in synaptic neuropil. A triangle threshold was applied to the pAMPK channel after Gaussian smoothing to 1.5um to define the cortex+neuropil region. Brp localization was used to define synaptic neuropil regions by first smoothing the Brp channel to create a mask that separated cortex from neuropil in ImageJ (gaussian = 0.6um) followed by hole-filling and binarization to produce segmented masks. Segmentation was visually verified for each optic lobe to ensure faithful definitions of cortex to neuropil boundaries. Next, mean pAMPK intensity from the raw data was measured within these defined neuropil and cortex regions to define the neuropil:cortex ratio of pAMPK enrichment per optic lobe (with one optic lobe per brain).

### Statistics

All data are reported as mean +/- SEM. Statistical tests are reported in the figure legends. For comparing responses such as AUC ratio and t-peak in subcellular compartments, the Kruskal-Wallis test was used followed by Mann-Whitney U tests with Holm-Bonferroni correction across all comparisons (α = 0.05). Because compartments were sometimes imaged in separate windows to accommodate large neurons spanning the medulla and lobula, ROIs could not be matched to the same neuron across compartments and neurons were therefore treated as independent. For other imaging experiments, differences across imaging conditions or genotypes were assessed using Welch’s t-test for groups of two and Welch’s one-way ANOVA for more than two groups, followed by Welch’s t-test against the control with Holm-Bonferroni correction for multiple comparisons. For the RNAi screen, experimental genotypes were compared with the pooled control (control + control-RNAi).

## References

1. G. A. Dienel, Brain Glucose Metabolism: Integration of Energetics with Function. Physiol. Rev. 99, 949–1045 (2019).

2. J. S. McCasland, L. S. Hibbard, GABAergic Neurons in Barrel Cortex Show Strong, Whisker-Dependent Metabolic Activation during Normal Behavior. J. Neurosci. 17, 5509–5527 (1997).

3. R. Ackermann, D. Finch, T. Babb, J. Engel, Increased glucose metabolism during long-duration recurrent inhibition of hippocampal pyramidal cells. J. Neurosci. 4, 251–264 (1984).

4. G. Buzsáki, K. Kaila, M. Raichle, Inhibition and Brain Work. Neuron 56, 771–783 (2007).

5. M. C. McKenna, Glutamate Pays Its Own Way in Astrocytes. Front. Endocrinol. 4 (2013).

6. J. V. Andersen, The Glutamate/ GABA-Glutamine Cycle: Insights, Updates, and Advances. J. Neurochem. 169, e70029 (2025).

7. W. Meier-Ruge, C. Bertoni-Freddari, P. Iwangoff, Changes in Brain Glucose Metabolism as a Key to the Pathogenesis of Alzheimer’s Disease. Gerontology, 246–252 (2009).

8. A. Szutowicz, H. Bielarczyk, A. Jankowska-Kulawy, T. Pawełczyk, A. Ronowska, Acetyl-CoA the Key Factor for Survival or Death of Cholinergic Neurons in Course of Neurodegenerative Diseases. Neurochem. Res. 38, 1523–1542 (2013).

9. Y. Tang, H. Zong, H. Kwon, Y. Qiu, J. B. Pessin, L. Wu, K. A. Buddo, I. Boykov, C. A. Schmidt, C.-T. Lin, P. D. Neufer, G. J. Schwartz, I. J. Kurland, J. E. Pessin, TIGAR deficiency enhances skeletal muscle thermogenesis by increasing neuromuscular junction cholinergic signaling. eLife 11, e73360 (2022).

10. C. Pulido, M. S. Gentry, T. A. Ryan, Neuromodulator control of energy reserves in dopaminergic neurons. [Preprint] (2025). 10.1101/2025.08.19.671144.

11. C. Pulido, T. A. Ryan, Synaptic vesicle pools are a major hidden resting metabolic burden of nerve terminals. Sci. Adv. 7, eabi9027 (2021).

12. C. Sobieski, M. J. Fitzpatrick, S. J. Mennerick, Differential Presynaptic ATP Supply for Basal and High-Demand Transmission. J. Neurosci. 37, 1888–1899 (2017).

13. V. Rangaraju, N. Calloway, T. A. Ryan, Activity-Driven Local ATP Synthesis Is Required for Synaptic Function. Cell 156, 825–835 (2014).

14. Q. Gan, S. Watanabe, The Effect of Acute ATP Depletion on Synaptic Vesicle Endocytosis at the Ultrastructural Level. Biophys. J. 116, 93a (2019).

15. D. Pathak, L. Y. Shields, B. A. Mendelsohn, D. Haddad, W. Lin, A. A. Gerencser, H. Kim, M. D. Brand, R. H. Edwards, K. Nakamura, The Role of Mitochondrially Derived ATP in Synaptic Vesicle Recycling. J. Biol. Chem. 290, 22325–22336 (2015).

16. C. M. Díaz-García, D. J. Meyer, N. Nathwani, M. Rahman, J. R. Martínez-François, G. Yellen, The distinct roles of calcium in rapid control of neuronal glycolysis and the tricarboxylic acid cycle. eLife 10, e64821 (2021).

17. D. J. Meyer, C. M. Díaz-García, N. Nathwani, M. Rahman, G. Yellen, The Na+/K+ pump dominates control of glycolysis in hippocampal dentate granule cells. eLife 11, e81645 (2022).

18. C. J. Groten, B. A. MacVicar, Mitochondrial Ca2+ uptake by the MCU facilitates pyramidal neuron excitability and metabolism during action potential firing. *Commun*. Biol. 5, 900 (2022).

19. I. Llorente-Folch, C. B. Rueda, B. Pardo, G. Szabadkai, M. R. Duchen, J. Satrustegui, The regulation of neuronal mitochondrial metabolism by calcium. J. Physiol. 593, 3447–3462 (2015).

20. G. Ashrafi, J. De Juan-Sanz, R. J. Farrell, T. A. Ryan, Molecular Tuning of the Axonal Mitochondrial Ca2+ Uniporter Ensures Metabolic Flexibility of Neurotransmission. Neuron 105, 678–687.e5 (2020).

21. A. K. Chouhan, M. V. Ivannikov, Z. Lu, M. Sugimori, R. R. Llinas, G. T. Macleod, Cytosolic Calcium Coordinates Mitochondrial Energy Metabolism with Presynaptic Activity. J. Neurosci. 32, 1233–1243 (2012).

22. E. J. Donovan, A. Agrawal, N. Liberman, J. I. Kalai, A. J. Adler, A. M. Lamper, H. Q. Wang, N. J. Chua, E. F. Koslover, E. L. Barnhart, Dendrite architecture determines mitochondrial distribution patterns in vivo. Cell Rep. 43, 114190 (2024).

23. M. C. Faits, C. Zhang, F. Soto, D. Kerschensteiner, Dendritic mitochondria reach stable positions during circuit development. eLife 5, e11583 (2016).

24. K. D. Longden, P. K. Rivlin, M. Januszewski, E. Neace, L. K. Scheffer, C. Ordish, J. Clements, E. Phillips, N. Smith, S. Takemura, L. Umayam, C. Walsh, E. A. Yakal, T. F. Musial, M. J. Devine, M. B. Reiser, S. M. Plaza, S. Berg, Connectomic Analysis of Mitochondria in the Central Brain of *Drosophila*. [Preprint] (2024). 10.1101/2024.04.21.590464.

25. G. Sager, P. Pfeiffer, H. Wu, F. Pallasdies, R. Gowers, S. Ravikumar, E. Wu, D. Colón-Ramos, S. Schreiber, D. A. Clark, Spatial and morphological organization of mitochondria in neurons across a connectome. Science 391, eads6674 (2026).

26. R. S. Stowers, L. J. Megeath, J. Górska-Andrzejak, I. A. Meinertzhagen, T. L. Schwarz, Axonal Transport of Mitochondria to Synapses Depends on Milton, a Novel Drosophila Protein. Neuron 36, 1063–1077 (2002).

27. E. E. Glater, L. J. Megeath, R. S. Stowers, T. L. Schwarz, Axonal transport of mitochondria requires milton to recruit kinesin heavy chain and is light chain independent. J. Cell Biol. 173, 545–557 (2006).

28. X. Wang, T. L. Schwarz, The Mechanism of Ca2+-Dependent Regulation of Kinesin-Mediated Mitochondrial Motility. Cell 136, 163–174 (2009).

29. X. Guo, G. T. Macleod, A. Wellington, F. Hu, S. Panchumarthi, M. Schoenfield, L. Marin, M. P. Charlton, H. L. Atwood, K. E. Zinsmaier, The GTPase dMiro Is Required for Axonal Transport of Mitochondria to Drosophila Synapses. Neuron 47, 379–393 (2005).

30. S. Jang, J. C. Nelson, E. G. Bend, L. Rodríguez-Laureano, F. G. Tueros, L. Cartagenova, K. Underwood, E. M. Jorgensen, D. A. Colón-Ramos, Glycolytic Enzymes Localize to Synapses under Energy Stress to Support Synaptic Function. Neuron 90, 278–291 (2016).

31. S. Jang, Z. Xuan, R. C. Lagoy, L. M. Jawerth, I. J. Gonzalez, M. Singh, S. Prashad, H. S. Kim, A. Patel, D. R. Albrecht, A. A. Hyman, D. A. Colón-Ramos, Phosphofructokinase relocalizes into subcellular compartments with liquid-like properties in vivo. Biophys. J. 120, 1170–1186 (2021).

32. S. B. Yu, H. Wang, R. G. Sanchez, N. M. Carlson, K. Nguyen, A. Zhang, Z. D. Papich, A. A. Abushawish, Z. Whiddon, W. Matysik, J. Zhang, T. C. Whisenant, M. Ghassemian, J. N. Koberstein, M. L. Stewart, S. A. Myers, G. Pekkurnaz, Neuronal activity-driven O-GlcNAcylation promotes mitochondrial plasticity. Dev. Cell 59, 2143–2157.e9 (2024).

33. H. Wang, J. W. Vant, A. Zhang, R. G. Sanchez, Y. Wu, M. L. Micou, V. Luczak, Z. Whiddon, N. M. Carlson, S. B. Yu, M. Jabbo, S. Yoon, A. A. Abushawish, M. Ghassemian, T. Masubuchi, Q. Gan, S. Watanabe, E. R. Griffis, M. Hammarlund, A. Singharoy, G. Pekkurnaz, Organization of a functional glycolytic metabolon on mitochondria for metabolic efficiency. Nat. Metab. 6, 1712–1735 (2024).

34. M.-V. Hinckelmann, A. Virlogeux, C. Niehage, C. Poujol, D. Choquet, B. Hoflack, D. Zala, F. Saudou, Self-propelling vesicles define glycolysis as the minimal energy machinery for neuronal transport. Nat. Commun. 7, 13233 (2016).

35. K. Takeda, T. Ueda, Effective Mechanism for Synthesis of Neurotransmitter Glutamate and its Loading into Synaptic Vesicles. Neurochem. Res. 42, 64–76 (2017).

36. J. Burré, W. Volknandt, The synaptic vesicle proteome. J. Neurochem. 101, 1448–1462 (2007).

37. A. Ikemoto, D. G. Bole, T. Ueda, Glycolysis and Glutamate Accumulation into Synaptic Vesicles. J. Biol. Chem. 278, 5929–5940 (2003).

38. M. Schläfer, W. Volknandt, H. Zimmermann, Putative Synaptic Vesicle Nucleotide Transporter Identified as Glyceraldehyde-3-Phosphate Dehydrogenase. J. Neurochem. 63, 1924–1931 (1994).

39. A. D. Wolfe, J. N. Koberstein, C. B. Smith, M. L. Stewart, I. J. Gonzalez, M. Hammarlund, A. A. Hyman, P. J. S. Stork, R. H. Goodman, D. A. Colón-Ramos, Local and dynamic regulation of neuronal glycolysis in vivo. Proc. Natl. Acad. Sci. 121, e2314699121 (2024).

40. T. A. Currier, M. M. Pang, T. R. Clandinin, Visual processing in the fly, from photoreceptors to behavior. GENETICS 224, iyad064 (2023).

41. A. Trevisiol, A. S. Saab, U. Winkler, G. Marx, H. Imamura, W. Möbius, K. Kusch, K.-A. Nave, J. Hirrlinger, Monitoring ATP dynamics in electrically active white matter tracts. eLife 6, e24241 (2017).

42. K. Schoknecht, F. Baeza-Lehnert, J. Hirrlinger, J. P. Dreier, J. Eilers, Spreading depolarizations exhaust neuronal ATP in a model of cerebral ischemia. [Preprint] (2024). 10.1101/2024.07.30.605834.

43. H. H. Yang, F. St-Pierre, X. Sun, X. Ding, M. Z. Lin, T. R. Clandinin, Subcellular Imaging of Voltage and Calcium Signals Reveals Neural Processing In Vivo. Cell 166, 245–257 (2016).

44. N. J. Allen, R. Káradóttir, D. Attwell, A Preferential Role for Glycolysis in Preventing the Anoxic Depolarization of Rat Hippocampal Area CA1 Pyramidal Cells. J. Neurosci. 25, 848–859 (2005).

45. J. P. Dreier, The role of spreading depression, spreading depolarization and spreading ischemia in neurological disease. Nat. Med. 17, 439–447 (2011).

46. J. N. Koberstein, M. L. Stewart, C. B. Smith, A. I. Tarasov, F. M. Ashcroft, P. J. S. Stork, R. H. Goodman, Monitoring glycolytic dynamics in single cells using a fluorescent biosensor for fructose 1,6-bisphosphate. Proc. Natl. Acad. Sci. 119, e2204407119 (2022).

47. M. Singh, A. D. Wolfe, A. A. Vishwanath, A. Tsives, I. J. Gonzalez, S. E. Emerson, R. Goodman, D. Colón-Ramos, Glycogen supports glycolytic plasticity in neurons. Proc. Natl. Acad. Sci. 122, e2509003122 (2025).

48. Z. Koshenov, A. C. Kokotos, L. Benedetti, J. L. Schwartz, T. A. Ryan, A genetic driver of epileptic encephalopathy impairs gating of synaptic glycolysis. [Preprint] (2025). 10.1101/2025.06.17.660213.

49. S. Bar, K. A. Wilson, T. A. U. Hilsabeck, S. Alderfer, E. B. Dammer, J. B. Burton, S. Shah, A. Holtz, E. M. Carrera, J. N. Beck, J. H. Chen, G. Kauwe, F. Seifar, A. Shantaraman, T. E. Tracy, N. T. Seyfried, B. Schilling, L. M. Ellerby, P. Kapahi, Neuronal glycogen breakdown mitigates tauopathy via pentose-phosphate-pathway-mediated oxidative stress reduction. Nat. Metab. 7, 1375–1391 (2025).

50. E. De Tredern, Y. Rabah, L. Pasquer, J. Minatchy, P.-Y. Plaçais, T. Preat, Glial glucose fuels the neuronal pentose phosphate pathway for long-term memory. Cell Rep. 36, 109620 (2021).

51. J. Tyler, A. A. Vishwanath, T. Menon, T. Duarah, R. Adhikari, J. N. Koberstein, D. Feliciano, I. Espinosa-Medina, D. Colón-Ramos, A. G. Tebo, Improved sensors for fructose-1,6-bisphosphate enable in vivo imaging of glycolysis. [Preprint] (2026). 10.64898/2026.04.29.721630.

52. J. Myeong, M. I. Stunault, V. A. Klyachko, G. Ashrafi, Metabolic Regulation of Single Synaptic Vesicle Exo- and Endocytosis in Hippocampal Synapses. [Preprint] (2023). 10.1101/2023.11.08.566236.

53. S. Jäger, C. Handschin, J. St.-Pierre, B. M. Spiegelman, AMP-activated protein kinase (AMPK) action in skeletal muscle via direct phosphorylation of PGC-1α. Proc. Natl. Acad. Sci. 104, 12017–12022 (2007).

54. D. G. Hardie, F. A. Ross, S. A. Hawley, AMPK: a nutrient and energy sensor that maintains energy homeostasis. Nat. Rev. Mol. Cell Biol. 13, 251–262 (2012).

55. C. F. Bringas, M. S. Ahangar, J. Cuenco, H. Liu, A. B. Addinsall, M. Lindahl, A. J. Ovens, M. A. Febbraio, M. Foretz, O. Göransson, J. W. Scott, E. Zeqiraj, K. Sakamoto, Mechanism and cellular actions of the potent AMPK inhibitor BAY-3827. Sci. Adv. 11, eadx2434 (2025).

56. S. Li, Z.-H. Sheng, Energy matters: presynaptic metabolism and the maintenance of synaptic transmission. Nat. Rev. Neurosci. 23, 4–22 (2022).

57. A. Nern, F. Loesche, S. Takemura, L. E. Burnett, M. Dreher, E. Gruntman, J. Hoeller, G. B. Huang, M. Januszewski, N. C. Klapoetke, S. Koskela, K. D. Longden, Z. Lu, S. Preibisch, W. Qiu, E. M. Rogers, P. Seenivasan, A. Zhao, J. Bogovic, B. S. Canino, J. Clements, M. Cook, S. Finley-May, M. A. Flynn, I. Hameed, A. M. C. Fragniere, K. J. Hayworth, G. P. Hopkins, P. M. Hubbard, W. T. Katz, J. Kovalyak, S. A. Lauchie, M. Leonard, A. Lohff, C. A. Maldonado, C. Mooney, N. Okeoma, D. J. Olbris, C. Ordish, T. Paterson, E. M. Phillips, T. Pietzsch, J. R. Salinas, P. K. Rivlin, P. Schlegel, A. L. Scott, L. A. Scuderi, S. Takemura, I. Talebi, A. Thomson, E. T. Trautman, L. Umayam, C. Walsh, J. J. Walsh, C. S. Xu, E. A. Yakal, T. Yang, T. Zhao, J. Funke, R. George, H. F. Hess, G. S. X. E. Jefferis, C. Knecht, W. Korff, S. M. Plaza, S. Romani, S. Saalfeld, L. K. Scheffer, S. Berg, G. M. Rubin, M. B. Reiser, Connectome-driven neural inventory of a complete visual system. Nature 641, 1225–1237 (2025).

58. S. Berg, I. R. Beckett, M. Costa, P. Schlegel, M. Januszewski, E. C. Marin, A. Nern, S. Preibisch, W. Qiu, S. Takemura, A. M. C. Fragniere, A. S. Champion, D.-Y. Adjavon, M. Cook, M. Gkantia, K. J. Hayworth, G. B. Huang, W. T. Katz, F. Kämpf, Z. Lu, C. Ordish, T. Paterson, T. Stürner, E. T. Trautman, C. R. Whittle, L. E. Burnett, J. Hoeller, F. Li, F. Loesche, B. J. Morris, T. Pietzsch, M. W. Pleijzier, V. Silva, Y. Yin, I. Ali, G. Badalamente, A. S. Bates, R. J. Beresford, J. Bogovic, P. Brooks, S. Cachero, B. S. Canino, B. Chaisrisawatsuk, J. Clements, A. Crowe, I. De Haan Vicente, G. Dempsey, E. Donà, M. Dos Santos, M. Dreher, C. R. Dunne, K. Eichler, S. Finley-May, M. A. Flynn, I. Hameed, G. P. Hopkins, P. M. Hubbard, L. Kiassat, J. Kovalyak, S. A. Lauchie, M. Leonard, A. Lohff, K. D. Longden, C. A. Maldonado, I. Moitra, S. S. Moon, C. Mooney, E. J. Munnelly, N. Okeoma, D. J. Olbris, A. Pai, B. Patel, E. M. Phillips, S. M. Plaza, A. Richards, J. Rivas Salinas, R. J. V. Roberts, E. M. Rogers, A. L. Scott, L. A. Scuderi, P. Seenivasan, L. Serratosa Capdevila, C. Smith, R. Svirskas, S. Takemura, I. Tastekin, A. Thomson, L. Umayam, J. J. Walsh, H. Whittome, C. S. Xu, E. A. Yakal, T. Yang, A. Zhao, R. George, V. Jain, V. Jayaraman, W. Korff, G. W. Meissner, S. Romani, J. Funke, C. Knecht, S. Saalfeld, L. K. Scheffer, S. Waddell, G. M. Card, C. Ribeiro, M. B. Reiser, H. F. Hess, G. M. Rubin, G. S. X. E. Jefferis, Sexual dimorphism in the complete Drosophila male central nervous system connectome. Cell 189, 5504–5526.e15 (2026).

59. R. Birks, F. C. MacIntosh, ACETYLCHOLINE METABOLISM OF A SYMPATHETIC GANGLION. Can. J. Biochem. Physiol. 39, 787–827 (1961).

60. G. E. Gibson, C. Peterson, Acetylcholine and oxidative metabolism in septum and hippocampus in vitro. J. Biol. Chem. 258, 1142–1145 (1983).

61. P. M. Borden, P. Zhang, A. V. Shivange, J. S. Marvin, J. Cichon, C. Dan, K. Podgorski, A. Figueiredo, O. Novak, M. Tanimoto, E. Shigetomi, M. A. Lobas, H. Kim, P. Zhu, Y. Zhang, W. S. Zheng, C. Fan, G. Wang, B. Xiang, L. Gan, G.-X. Zhang, K. Guo, L. Lin, Y. Cai, A. G. Yee, A. Aggarwal, H. Bao, X. Lou, E. R. Chapman, C. P. Ford, D. Rees, D. Dietrich, B. S. Khakh, J. S. Dittman, W.-B. Gan, M. Koyama, V. Jayaraman, J. F. Cheer, H. A. Lester, J. J. Zhu, L. Looger, A Fast Genetically Encoded Fluorescent Sensor for Faithful& *in vivo*&Acetylcholine Detection in Mice, Fish, Worms and Flies. SSRN Electron. J., doi: 10.2139/ssrn.3554080 (2020).

62. J. S. Marvin, B. Scholl, D. E. Wilson, K. Podgorski, A. Kazemipour, J. A. Müller, S. Schoch, F. J. U. Quiroz, N. Rebola, H. Bao, J. P. Little, A. N. Tkachuk, E. Cai, A. W. Hantman, S. S.-H. Wang, V. J. DePiero, B. G. Borghuis, E. R. Chapman, D. Dietrich, D. A. DiGregorio, D. Fitzpatrick, L. L. Looger, Stability, affinity, and chromatic variants of the glutamate sensor iGluSnFR. Nat. Methods 15, 936–939 (2018).

63. A. Szutowicz, M. Tomaszewicz, A. Jankowska, B. Madziar, H. Bielarczyk, Acetyl-CoA metabolism in cholinergic neurons and their susceptibility to neurotoxic inputs. Metab. Brain Dis. 15, 29–44 (2000).

64. E. A. Newsholme, P. H. Sugden, T. Williams, Effect of citrate on the activities of 6-phosphofructokinase from nervous and muscle tissues from different animals and its relationships to the regulation of glycolysis. Biochem. J. 166, 123–129 (1977).

65. B. A. Webb, A. M. Dosey, T. Wittmann, J. M. Kollman, D. L. Barber, The glycolytic enzyme phosphofructokinase-1 assembles into filaments. J. Cell Biol. 216, 2305–2313 (2017).

66. A. H. Romano, T. Conway, Evolution of carbohydrate metabolic pathways. Res. Microbiol. 147, 448–455 (1996).

67. D. Yu, Z. Shen, P. Wu, X. Guan, T. Chen, Y. Jin, Z. Hu, L. Ni, F. Wang, J. Chen, L. Long, HFS-Triggered AMPK Activation Phosphorylates GSK 3β and Induces E-LTP in Rat Hippocampus *In Vivo*. CNS Neurosci. Ther. 22, 525–531 (2016).

68. G. Ashrafi, Z. Wu, R. J. Farrell, T. A. Ryan, GLUT4 Mobilization Supports Energetic Demands of Active Synapses. Neuron 93, 606–615.e3 (2017).

69. Y. Z. Kurmangaliyev, J. Yoo, J. Valdes-Aleman, P. Sanfilippo, S. L. Zipursky, Transcriptional Programs of Circuit Assembly in the Drosophila Visual System. Neuron 108, 1045–1057.e6 (2020).

70. S. A. Hawley, J. Boudeau, J. L. Reid, K. J. Mustard, L. Udd, T. P. Mäkelä, D. R. Alessi, D. G. Hardie, Complexes between the LKB1 tumor suppressor, STRADα/β and MO25α/β are upstream kinases in the AMP-activated protein kinase cascade. J. Biol. 2, 28 (2003).

71. A. Woods, S. R. Johnstone, K. Dickerson, F. C. Leiper, L. G. D. Fryer, D. Neumann, U. Schlattner, T. Wallimann, M. Carlson, D. Carling, LKB1 Is the Upstream Kinase in the AMP-Activated Protein Kinase Cascade. Curr. Biol. 13, 2004–2008 (2003).

72. E. Van Schaftingen, M. F. Jett, L. Hue, H. G. Hers, Control of liver 6-phosphofructokinase by fructose 2,6-bisphosphate and other effectors. Proc. Natl. Acad. Sci. 78, 3483–3486 (1981).

73. R. S. Jope, D. J. Jenden, The Utilization of Choline and Acetyl Coenzyme A for the Synthesis of Acetylcholine. J. Neurochem. 35, 318–325 (1980).

74. M. Sarter, V. Parikh, Choline transporters, cholinergic transmission and cognition. Nat. Rev. Neurosci. 6, 48–56 (2005).

75. P. C. Rosen, P. Fu, B. Ferrán, E. Kim, D. J. Brooks, D. C. Lim, C. M. Díaz-García, G. Yellen, Activity-dependent citrate dynamics in neurons. Proc. Natl. Acad. Sci. 122, e2519902122 (2025).

76. E. C. Ballinger, M. Ananth, D. A. Talmage, L. W. Role, Basal Forebrain Cholinergic Circuits and Signaling in Cognition and Cognitive Decline. Neuron 91, 1199–1218 (2016).

77. G. M. Kerbler, J. Fripp, C. C. Rowe, V. L. Villemagne, O. Salvado, S. Rose, E. J. Coulson, Basal forebrain atrophy correlates with amyloid β burden in Alzheimer’s disease. NeuroImage Clin. 7, 105–113 (2015).

78. M. Ashrafi, L. Fraticelli, G. Castrillón, V. Riedl, Metabolism-weighted brain connectome reveals synaptic integration and vulnerability to neurodegeneration. Proc. Natl. Acad. Sci. 123, e2531706123 (2026).

79. S. M. Plaza, J. Clements, T. Dolafi, L. Umayam, N. N. Neubarth, L. K. Scheffer, S. Berg, neuPrint: An open access tool for EM connectomics. *Front*. Neuroinformatics 16, 896292 (2022).

