## Supplementary Materials for "Neuronal activity regulates glycolysis to sculpt synaptic transmission"

### Materials and Methods

Materials

| REAGENT or RESOURCE | SOURCE | IDENTIFIER |
|---|---|---|
| **Antibodies** | | |
| Nc82 (anti-mouse IgG1), 1:10 | Developmental Studies Hybridoma Bank | RRID: AB_2314866 |
| phospho-AMPK (anti-rabbit), 1:500 | Cell Signaling Technology | Cat# #2535, RRID: AB_331250 |
| Goat anti-Mouse IgG1 Cross-Adsorbed Secondary Antibody, Alexa Fluor 488, 1:500 | Thermofisher scientific | RRID: AB_2535764 |
| Goat anti-Rabbit Cross-Adsorbed Secondary Antibody, Cyanine3, 1:500 | Thermofisher scientific | RRID: AB_10563288 |
| **Pharmacology** | | |
| Sodium azide ($NaN^3$) | Sigma-Aldrich | Cat # 71289 |
| Dorsomorphin (compound C) | Sigma-Aldrich | Cat #171264 |
| BAY-3827 | Sigma-Aldrich | Cat # SML3795 |
| **Experimental models: Organisms/strains** | | |
| *D. melanogaster*: Tm1-Gal4: +*; +; R74G01-Gal4* | Janelia FlyLight Project | RRID: BDSC_ 39868 |
| *D. melanogaster*: L1-split-Gal4: w[1118]; *VT027316-p65ADZp (attP40); R40F12-GAL4-ZpGdbd (attP2)* | Obtained from G. Rubin ( 10.1016/j.cub.2019.10.035) | RRID: BDSC_601133 and JRC_SS03734 |
| *D. melanogaster*: L2-split-Gal4: $w^{1118}$; *R53G02-p65ADZp(attP40); R29G11-ZpGdbd (attP2)* | Janelia Research Campus | JRC_SS00690 |
| *D. melanogaster*: Tm2-split-Gal4: $w^{1118}$; *R26E12-p65ADZp(attP40); R48D11-ZpGdbd (attP2)* | Janelia Research Campus | JRC_SS00048 |
| *D. melanogaster*: Tm3-split-Gal4: *w1118; R13E12-p65ADZp(attP40); R59C10-ZpGdbd(attP2)* | Janelia Research Campus | JRC_SS00313 |
| *D. melanogaster*: Tm9-split-Gal4 | Janelia Research Campus | JRC_SS00307 |
| *D. melanogaster*: Mi1-split-Gal4: $w^{1118}$; *R55C05-p65ADZp(attP40); R71D01-ZpGdbd (attP2)* | Janelia Research Campus | JRC_SS00955 |
| *D. melanogaster*: Mi9-split-Gal4: *w1118; R48A07-p65ADZp(attP40); VT046779-ZpGdbd(attP2)* | Janelia Research Campus | JRC_SS02432 |
| *D. melanogaster*: {20xUAS-Hylight}attp5 | This study | |
| *D. melanogaster*: {20xUAS-HYlight2}attp5 | This study | |

| | | |
|---|---|---|
| *D. melanogaster:* P{20xUAS-SF-iGluSnFR.A184V}attP40 | This study | |
| *D. melanogaster:* P{20xUAS-iAChSnFR (V188P)}attP40 | This study | |
| *D. melanogaster*: {20xUAS-IVS-NES-jRGECO1a-p10}VK00005 | Bloomington Drosophila Resource Center | RRID:BDSC_63794 |
| **RNAi lines** | | |
| *D. melanogaster*: UAS-TNT | Bloomington Drosophila Resource Center | RRID:BDSC_28838 |
| *D. melanogaster*: control-RNAi: | Bloomington Drosophila Resource Center | RRID:BDSC_36303 |
| *D. melanogaster*: pfk-RNAi: | Bloomington Drosophila Resource Center | RRID:BDSC_34336 |
| *D. melanogaster*: pfk-OE: | Bloomington Drosophila Resource Center | RRID:BDSC_60675 |
| *D. melanogaster*: treh-RNAi: | Bloomington Drosophila Resource Center | RRID:VDRC_v30730 |
| *D. melanogaster*: glut1-RNAi: | Bloomington Drosophila Resource Center | RRID:BDSC_28645 |
| *D. melanogaster*: ald-RNAi: | Bloomington Drosophila Resource Center | RRID:BDSC_26301 |
| *D. melanogaster*: gapdh3-RNAi: | Bloomington Drosophila Resource Center | RRID:BDSC_62213 |
| *D. melanogaster*: pgk-RNAi 1: | Bloomington Drosophila Resource Center | RRID:BDSC_35220 |
| *D. melanogaster*: pgk-RNAi 2: | Bloomington Drosophila Resource Center | RRID:VDRC_v110081 |
| *D. melanogaster*: Eno-RNAi: | Bloomington Drosophila Resource Center | RRID:BDSC_26300 |
| *D. melanogaster*: PyK-RNAi: | Bloomington Drosophila Resource Center | RRID:VDRC_v35165 |
| *D. melanogaster*: glyp-RNAi: | Bloomington Drosophila Resource Center | RRID:BDSC_33634 |
| *D. melanogaster*: parkin-RNAi: | Vienna Drosophila Resource Center | RRID:VDRC_v47637 |

| | | |
|---|---|---|
| *D. melanogaster*: milt-RNAi: | Bloomington Drosophila Resource Center | RRID:BDSC_28385 |
| *D. melanogaster*: miro-RNAi: | Bloomington Drosophila Resource Center | RRID:BDSC_27695 |
| *D. melanogaster*: oga-RNAi: | Vienna Drosophila Resource Center | RRID:VDRC_v41822 |
| *D. melanogaster*: ogt-RNAi: | Vienna Drosophila Resource Center | RRID:VDRC_v18610 |
| *D. melanogaster*: ap2-RNAi: | Bloomington Drosophila Resource Center | RRID:BDSC_27322 |
| *D. melanogaster*: chc-RNAi: | Bloomington Drosophila Resource Center | RRID:BDSC_34742 |
| *D. melanogaster*: unc104-RNAi: | Bloomington Drosophila Resource Center | RRID:BDSC_43264 |
| *D. melanogaster*: vacht-RNAi: | Bloomington Drosophila Resource Center | RRID:BDSC_27684 |
| *D. melanogaster*: vhaAC45RP-RNAi: | Bloomington Drosophila Resource Center | RRID:BDSC_64549 |
| *D. melanogaster*: Pka-C1-RNAi: | Bloomington Drosophila Resource Center | RRID:BDSC_31277 |
| *D. melanogaster*: CaMKII-dsRNA: | Bloomington Drosophila Resource Center | RRID:BDSC_29401 |
| *D. melanogaster*: AMPK-RNAi 1: | Bloomington Drosophila Resource Center | RRID:BDSC_57785 |
| *D. melanogaster*: AMPK-RNAi 2: | Vienna Drosophila Resource Center | RRID:VDRC_v106200 |
| *D. melanogaster*: AMPK-OE: | Bloomington Drosophila Resource Center | RRID:BDSC_32108 |
| *D. melanogaster*: snf4a-RNAi: | Bloomington Drosophila Resource Center | RRID:BDSC_26291 |
| *D. melanogaster*: alc-RNAi: | Bloomington Drosophila Resource Center | RRID:BDSC_41606 |
| *D. melanogaster*: lkb-RNAi: | Bloomington Drosophila Resource Center | RRID:BDSC_35151 |

| | | |
|---|---|---|
| *D. melanogaster*: tak1-RNAi: | Bloomington Drosophila Resource Center | RRID:BDSC_33404 |
| *D. melanogaster*: mtor-RNAi: | Bloomington Drosophila Resource Center | RRID:BDSC_35578 |
| *D. melanogaster*: acly-RNAi: | Bloomington Drosophila Resource Center | RRID:BDSC_65175 |
| **Software and algorithms** | | |
| ImageJ (FIJI) | ImageJ | RRID: SCR_003070 |
| GraphPad Prism 10.2.2 | GraphPad | RRID: SCR_002798 |
| Python/Jupyter Labs (4.0.2) | Python | RRID: SCR_008394 |
| Visanalysis | Clandinin Lab | https://github.com/ClandininLab/visanalysis |
| Visprotocol | Clandinin Lab | https://github.com/ClandininLab/visprotocol |
| Original python code | This paper | https://github.com/theisenemma/theisen_analysis |

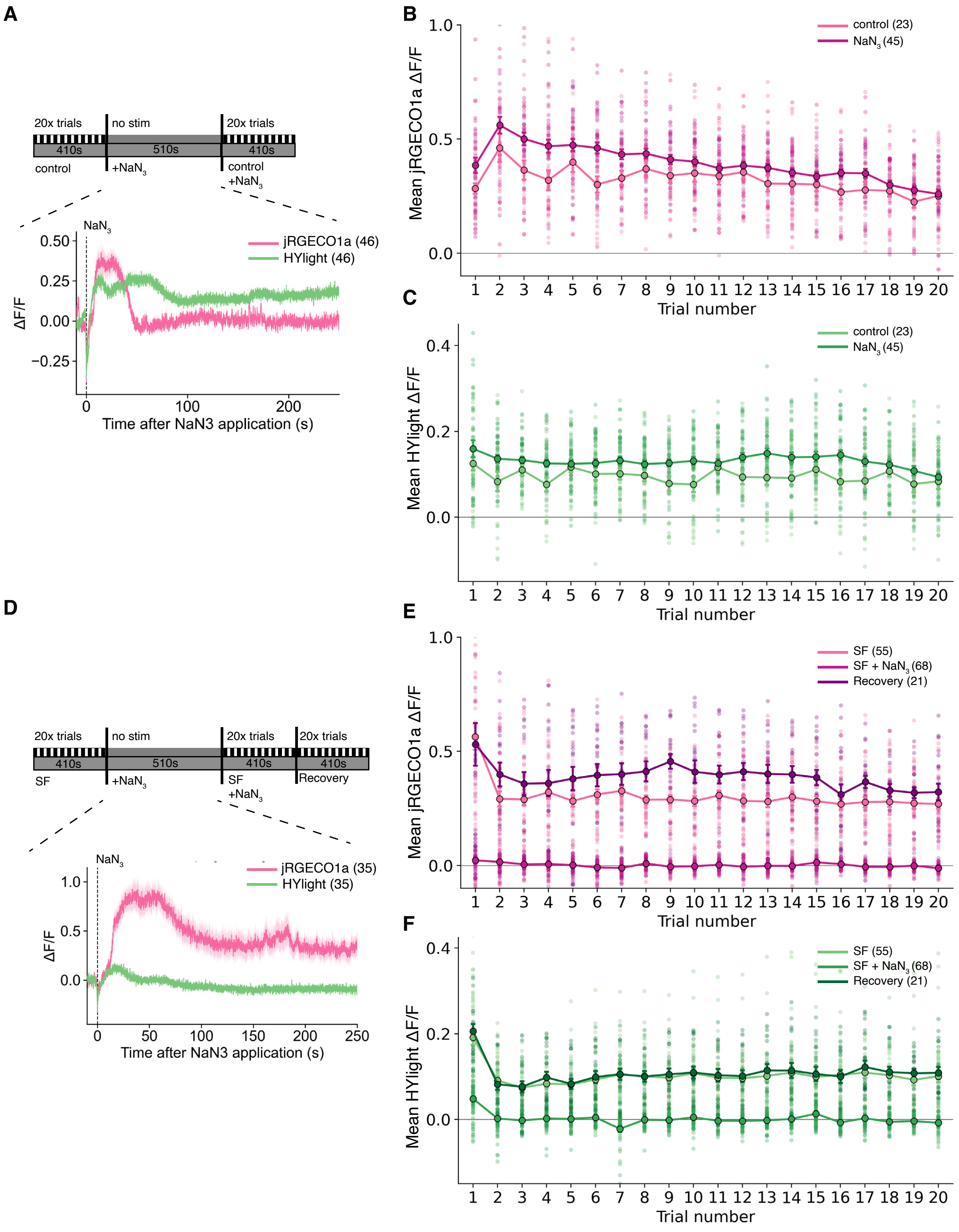
A
20x trials
no stim
20x trials
410s
510s
410s
control
+NaN$_3$
control
+NaN$_3$
NaN$_3$
jRGECO1a (46)
HYlight (46)
ΔF/F
Time after NaN3 application (s)
B
control (23)
NaN$_3$ (45)
Mean jRGECO1a ΔF/F
Trial number
C
control (23)
NaN$_3$ (45)
Mean HYlight ΔF/F
Trial number
D
20x trials
no stim
20x trials
20x trials
410s
510s
410s
410s
SF
+NaN$_3$
SF
+NaN$_3$
Recovery
NaN$_3$
jRGECO1a (35)
HYlight (35)
ΔF/F
Time after NaN3 application (s)
E
SF (55)
SF + NaN$_3$ (68)
Recovery (21)
Mean jRGECO1a ΔF/F
Trial number
F
SF (55)
SF + NaN$_3$ (68)
Recovery (21)
Mean HYlight ΔF/F
Trial number

**Fig. S1. The effect of sodium azide and sugar deprivation on activity and glycolysis.**

(**A**) Top: Imaging experiment schematic. Responses to visual stimuli were first recorded in control saline, followed by addition of NaN3 into the imaging saline to a final concentration of 2mM, during which stimulus-free responses were monitored for 510 seconds, followed by a final imaging trial. Middle: immediate responses to NaN3 in Tm1 neurons expressing jRGECO1a (magenta) and HYlight (green). Under these conditions, Tm1 transiently experiences a substantial increase in intracellular calcium concentration coincident with ATP depletion, before recovering after approximately 5-7 minutes. (**B**)Average jRGECO1a responses during the imaging series for each trial, before (magenta) and after (light pink) NaN3 addition.(**C**) Average HYlight responses during the imaging series for each trial, before (green) and after (light green) NaN3 addition. (**D**)Top: Imaging experiment schematic in SF saline. First, responses to visual stimuli were recorded, followed by injection of NaN3 and a subsequent recording of responses to visual stimuli, after which control (sugar-containing) saline was re-perfused onto the brain and neurons were reimaged while presenting visual stimuli. Middle: immediate responses to NaN3 in Tm1 neurons expressing jRGECO1a (magenta) and HYlight (green). (**E**) Average jRGECO1a responses during the imaging series for each trial, before (magenta) after (light pink) NaN3 addition, and after recovery saline (dark purple). (**F**)Average HYlight responses during the imaging series for each trial, before (green) and after (light green) NaN3 addition, and after recovery saline (dark green).

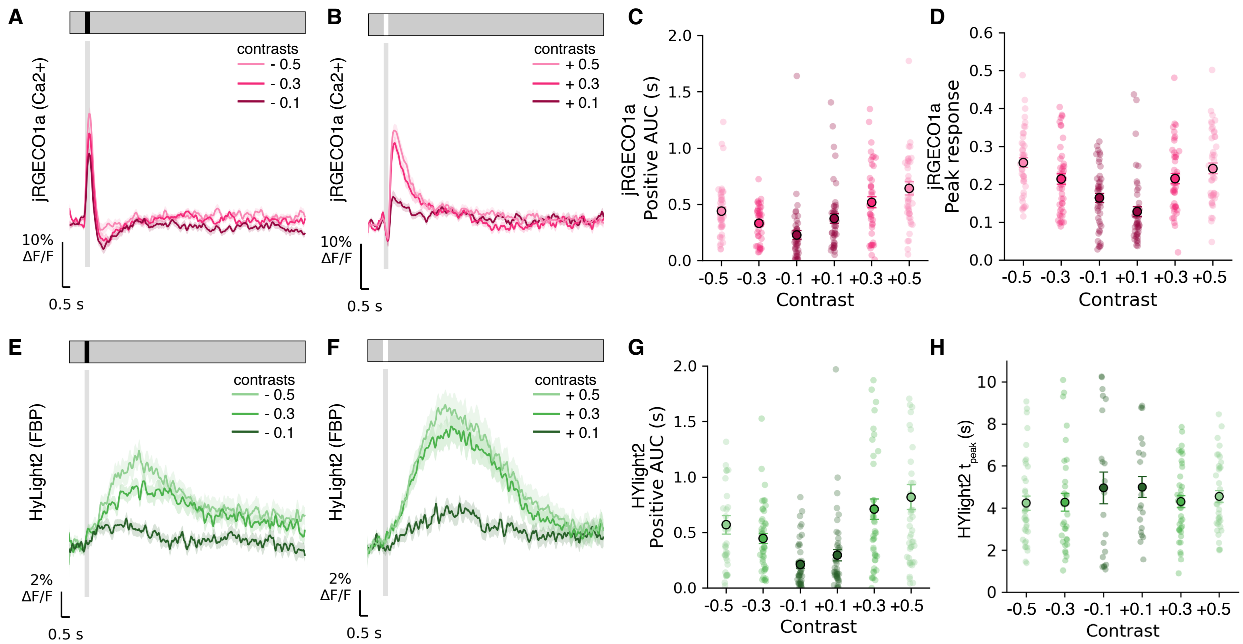


**Fig. S2. Neuronal activity induces glycolytic transients in response to 300ms- light flashes**
(**A**) Mean jRGECO1a responses collected during a 300ms dark flash stimulus across three dark contrasts and (**B**) three light contrasts. (**C**) Positive AUC of the jRGECO1a response across the contrast series. (**D**) Peak jRGECO1a response for each contrast. (**E**) HYlight2 responses collected during a 300ms dark flash stimulus across three dark contrasts. (**F**) HYlight2 responses collected during a 300ms light flash stimulus across three light contrasts. (**G**) Positive AUC of the HYlight2 response across six contrasts. (**H**) Time-to-peak of the HYlight2 response to contrast series.

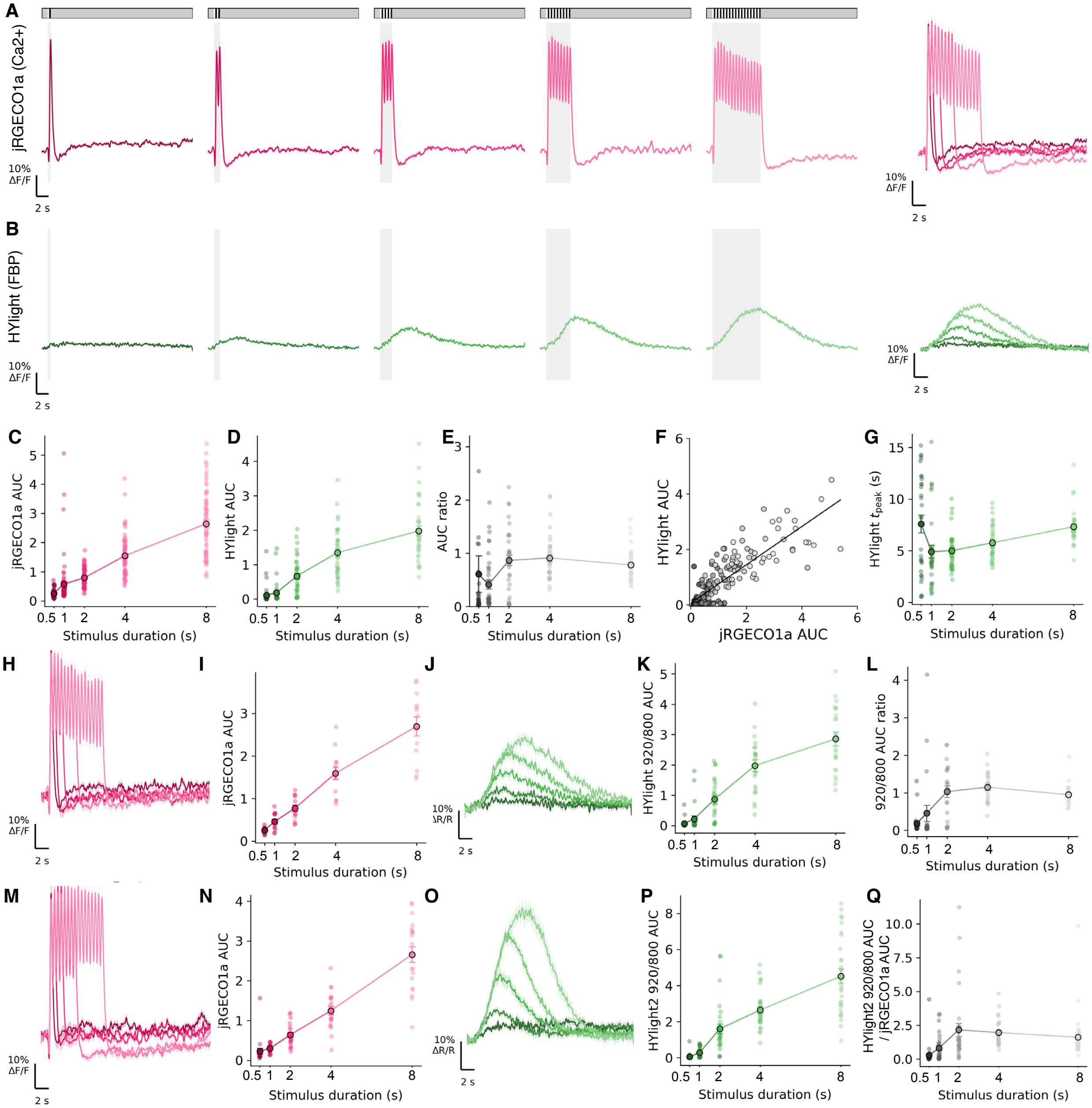


**Fig. S3. Quantifying ratiometric HYlight signals**

(**A**) Mean jRGECO1a responses to duration stimuli (**B**) Mean HYlight responses collected simultaneously with (**A**). (**C**) jRGECO1a AUC and (**D**) HYlight AUC and (**E)** HYlight ratio across stimulus durations. (**F**) HYlight AUC plotted against jRGECO1a AUC, with each dot representing a neuron (**G**) Time-to-peak of the HYlight response remains relatively stable across duration stimuli. (**H-L**) Ratiometric HYlight measurements obtained by sequentially imaging the FBP-unbound (800nm) and FBP-bound (920nm) responses of the sensor. (**H**) Mean jRGECO1a responses across stimulus durations (**I**) jRGECO1a AUC across the stimulus durations. (**J**) Mean ratiometric HYlight 920/800nm acquired simultaneously with (**H**). (**K**) Plotting the HYlight 920/800 AUC across the stimulus duration series. (**L**) The AUC ratio of HYlight 920/800 AUC normalized to the jRGECO1a AUC across stimulus durations. (**M-Q**) Same as in (**H-L**) but acquired using the HYlight2 sensor.

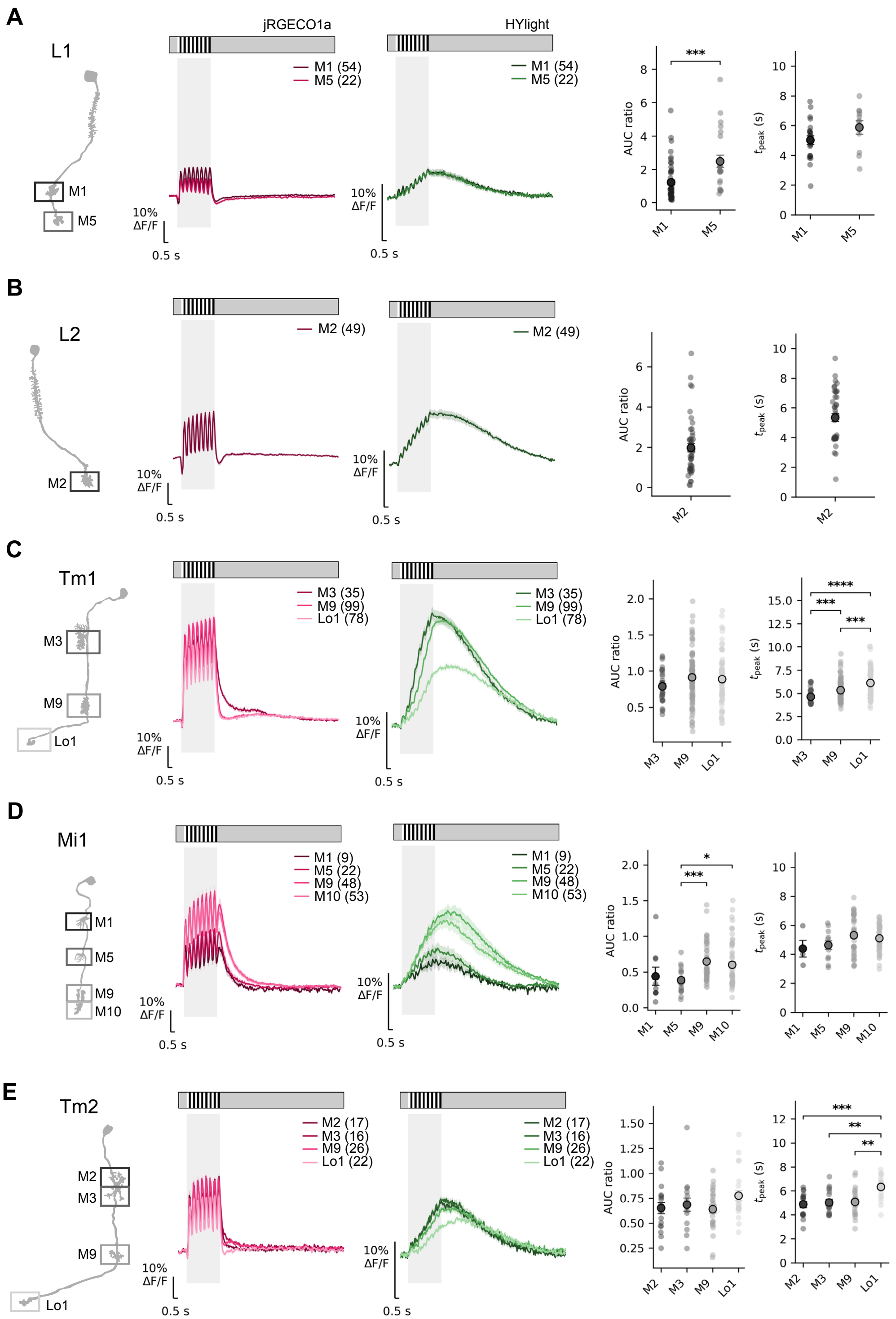
A
L1
jRGECO1a
HYlight
M1 (54)
M5 (22)
M1
M5
10% ΔF/F
0.5 s
AUC ratio
$t_{peak}$ (s)
***
B
L2
M2 (49)
M2
C
Tm1
M3 (35)
M9 (99)
Lo1 (78)
M3
M9
Lo1
****
***
D
Mi1
M1 (9)
M5 (22)
M9 (48)
M10 (53)
M1
M5
M9
M10
*
E
Tm2
M2 (17)
M3 (16)
M9 (26)
Lo1 (22)
M2
M3
**

**Fig. S4. Activity-induced glycolysis is flexible across subcellular compartments and neuron types**

(**A-E**) Glycolytic responses measured in (**A**) L1 (**B**) L2 (**C**) Tm1 (**D**) Mi1 and (**E**) Tm2 neurons. Left: schematic of neuron with compartments framed; Middle left = Mean jRGECO1a ($Ca^{2+}$) response of each compartment in neuron indicated. Middle right: Mean HYlight (FBP) response of each compartment in neuron indicated. Right = AUC ratio and HYlight tpeak per compartment. Each dot represents a neuron and circled dot indicates the mean. Error bars = SEM. Overall differences across compartments were assessed using Welch's one-way ANOVA. Experimental genotypes were compared with the pooled control (control + control-RNAi) using two-sided Welch's t-tests with Holm correction for multiple comparisons.
* $p < 0.05$, ** $p < 0.01$, *** $p < 0.001$, **** $p < 0.0001$.

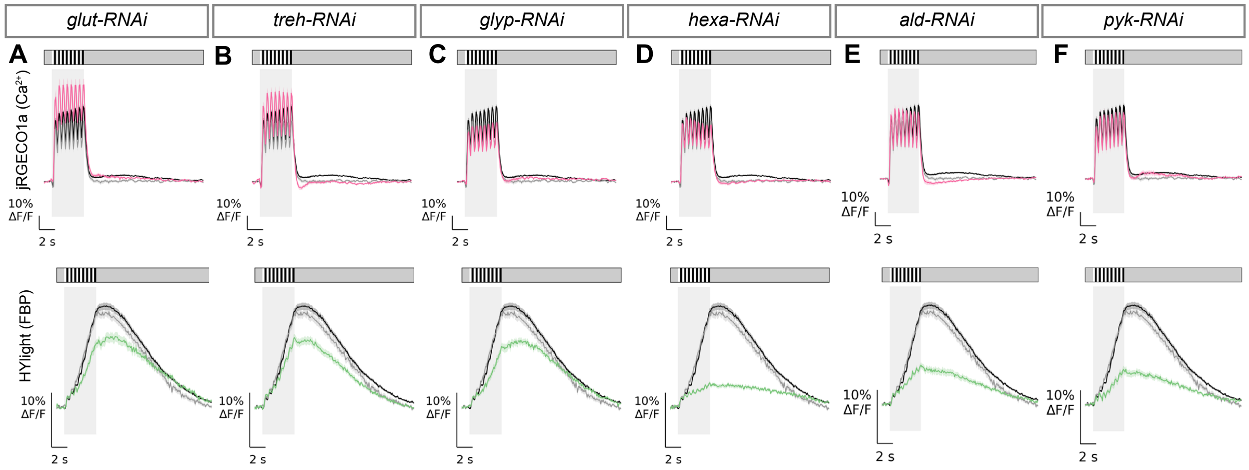


**Fig. S5. Activity-induced glycolysis consists of multiple essential enzymatic steps**
(**A-F**) Top: Mean jRGECO1a (Ca2+) responses plotted for control (black), control-RNAi (gray) and the corresponding RNAi condition (magenta) denoted at the top. Bottom: Mean HYlight (FBP) plotted for control (black), control-RNAi (gray) and the corresponding RNAi manipulation (green) labeled at the top. In (**A-G**), shading around each trace is SEM, the bar above indicates the visual stimulus, and the shaded region denotes the stimulus period. Scale bars, 10% ΔF/F and 2s.
* p <0.05, ** p <0.01, *** p <0.001, **** p<0.0001.

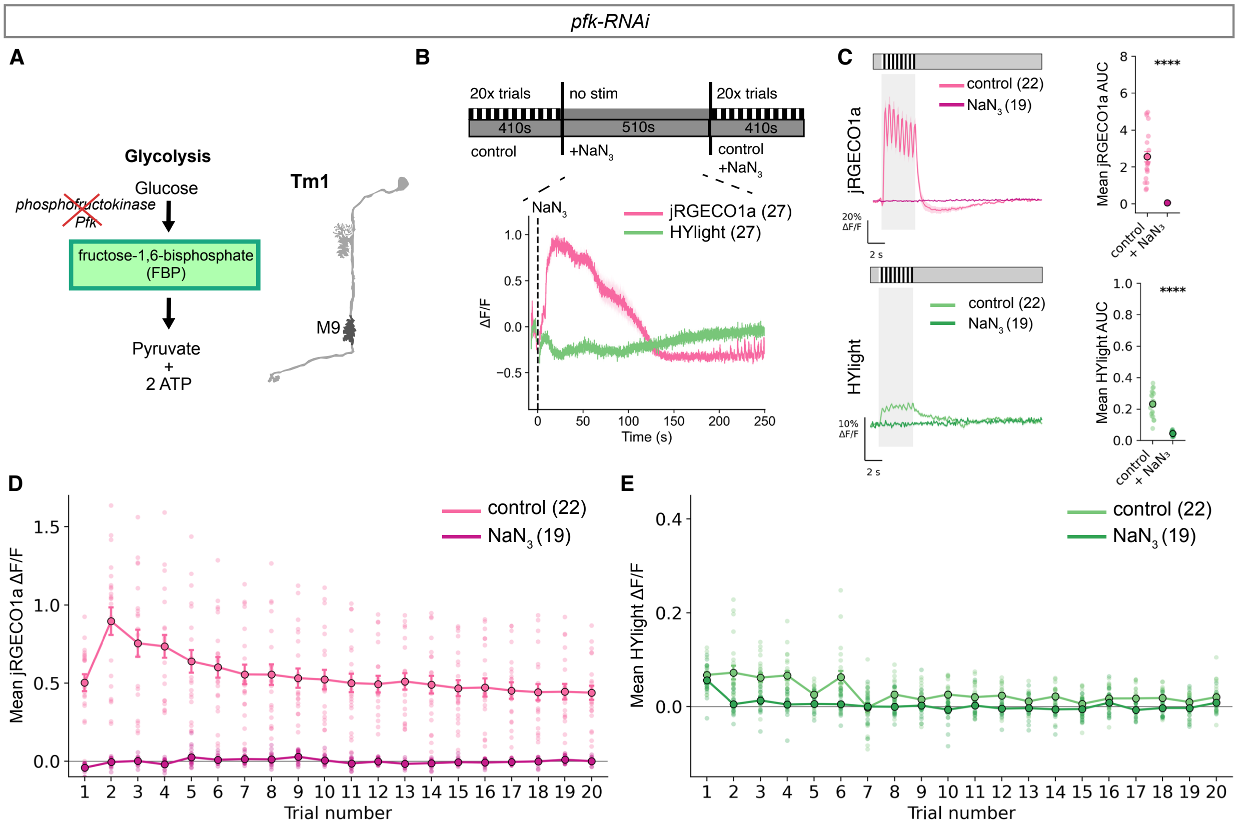


**Fig. S6. Glycolysis is required cell autonomously to sustain neuronal activity when oxphos is blocked.**

(**A-E**) All experiments were performed after pfk knockdown in Tm1 neurons (**A**) Schematic of RNAi-mediated knockdown of Pfk in Tm1 neurons. (**B**) Immediate jRGECO1a and HYlight responses to NaN3 application in Tm1 neurons with cell-type specific Pfk knockdown. Top schematic depicts the experiment setup: visual responses were collected before and after NaN3 application, and NaN3 was injected at the beginning of a 510-second long imaging window with no stimuli shown. (**C**) Top: mean jRGECO1a (Ca2+) responses to visual stimuli, before NaN3 application (control, light pink) and after (NaN3, magenta) which resulted in flatlined responses. Top right: AUC calculation of jRGECO1a response. Bottom: Mean HYlight (FBP) responses to visual stimuli, before NaN3 application (control, light green) and after (NaN3, green), which resulted in flatlined responses. Bottom right: AUC calculation of HYlight response. (**D**) Mean jRGECO1a max ΔF/F during each trial before (control, light pink) and after (NaN3, magenta) NaN3 appplication. Responses remained flatlined throughout the imaging series after NaN3 application, consistent with loss of signaling observed after NaN3 application in sugar-free saline of control neurons. (**E**) Mean HYlight (FBP) max ΔF/F for each imaging trial before (control, light green) and after (NaN3, green) NaN3 appplication. Responses remained flatlined throughout the imaging series after NaN3 application.

Overall differences across genotypes were assessed using Welch's one-way ANOVA. Experimental genotypes were compared with the pooled control (control + control-RNAi) using two-sided Welch's t-tests with Holm correction for multiple comparisons.
* $p < 0.05$, ** $p < 0.01$, *** $p < 0.001$, **** $p < 0.0001$.

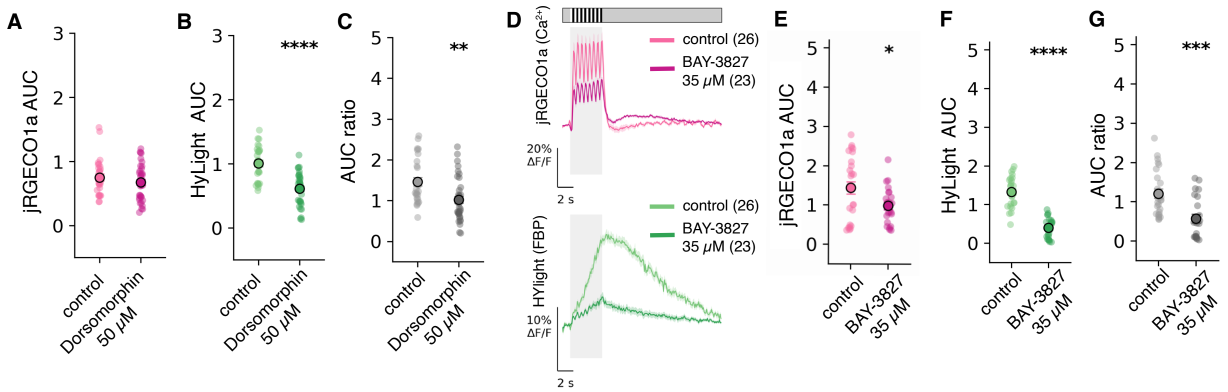


**Fig. S7. Pharmacological inhibition of AMPK disrupts activity-induced glycolytic responses** (**A-C**) Responses before and after 50uM Dorsomorphin application quantified as AUC. (**A**) jRGECO1a AUC before and after Dorsomorphin (**B**) HYlight AUC before and after Dorsomorphin (**B**) AUC ratio before and after Dorsomorphin (**D-G**) Responses before and after 35uM BAY-3827 application. (**D**) Top: Mean jRGECO1a responses, bottom: mean HYlight responses (**E**) jRGECO1a AUC before and after 35uM BAY-3827 application. (F) HYlight AUC before and after 35uM BAY-3827 application (**G**) AUC ratio before and after 35uM BAY-3827 application

* p <0.05, ** p <0.01, *** p <0.001, **** p<0.0001.

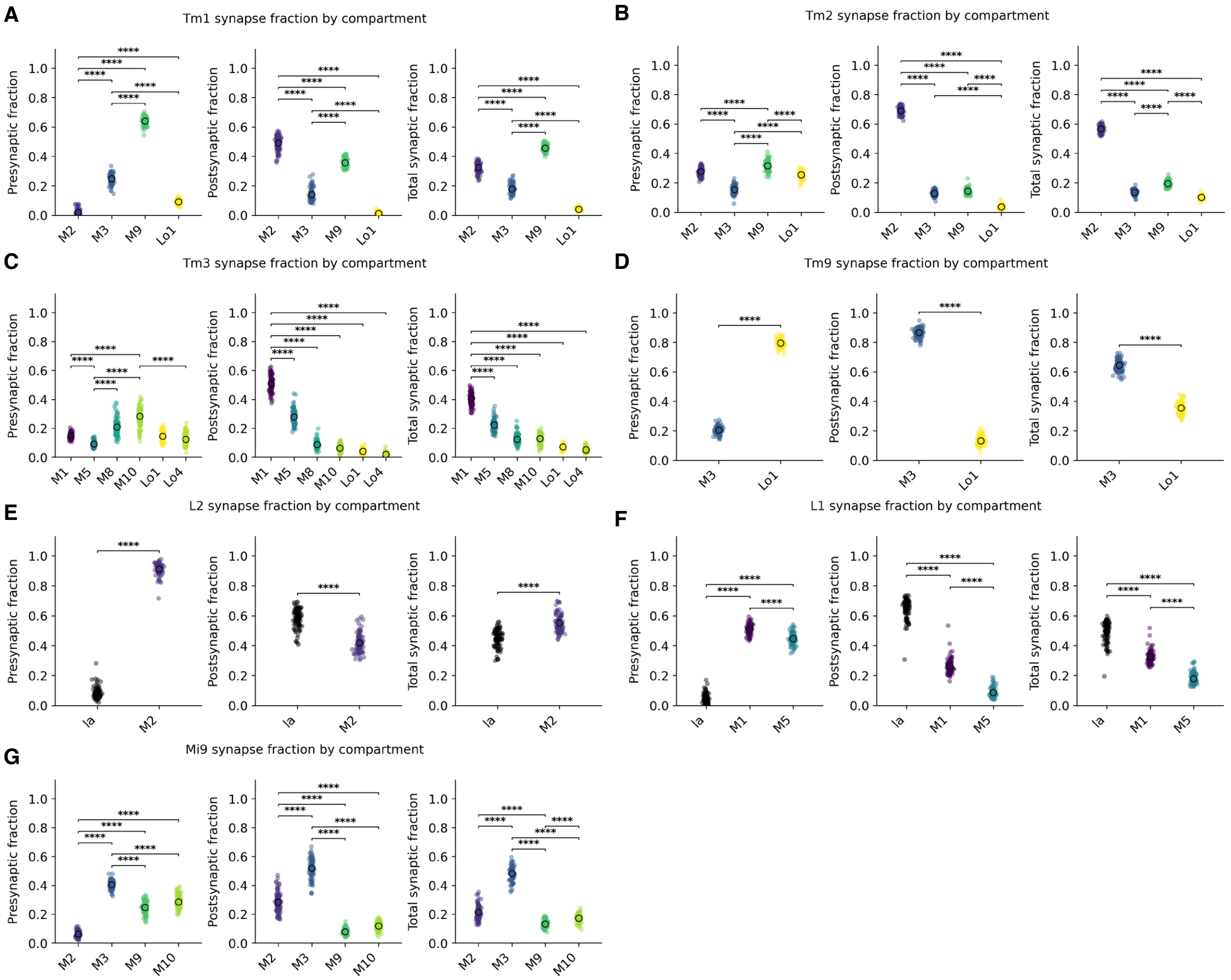


**Fig. S8. Connectome analysis of subcellular synapse distribution in medulla neurons**
(**A-G**) Synapse fractions by compartment for (**A**) Tm1, (**B**) Tm2, (**C**) Tm3, (**D**), Tm9 (**E**) L2 (**F**) L1 and (**G**) Mi9. Each neuron type is represented by 100 neurons, with the exception of L1 (91) and L2 (71) (see methods). Data obtained from the Optic Lobe v1.1 neuPrint connectome.

Synapse fractions across compartments were compared within neurons using the Friedman test (Wilcoxon signed-rank test for two compartments), followed by pairwise Wilcoxon signed-rank tests with Holm–Bonferroni correction ($\alpha = 0.05$).
* p <0.05, ** p <0.01, *** p <0.001, **** p<0.0001.

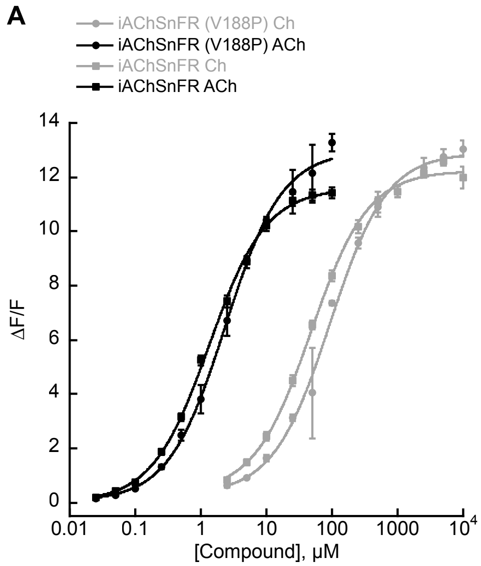


**Fig. S9. In vitro characterization of iAchSnFR (V188P)**

(**A**) In vitro titrations of purified WT iAchSnFR compared to the iAchSnFR (V188P) variant with acetylcholine (black lines) or choline (grey lines). The indicated concentrations of compound were added to 200nM of purified sensor in PBS and GFP fluorescence immediately measured using a fluorescence plate reader. Data plotted are mean ± S.D., n=3 replicate experiments.

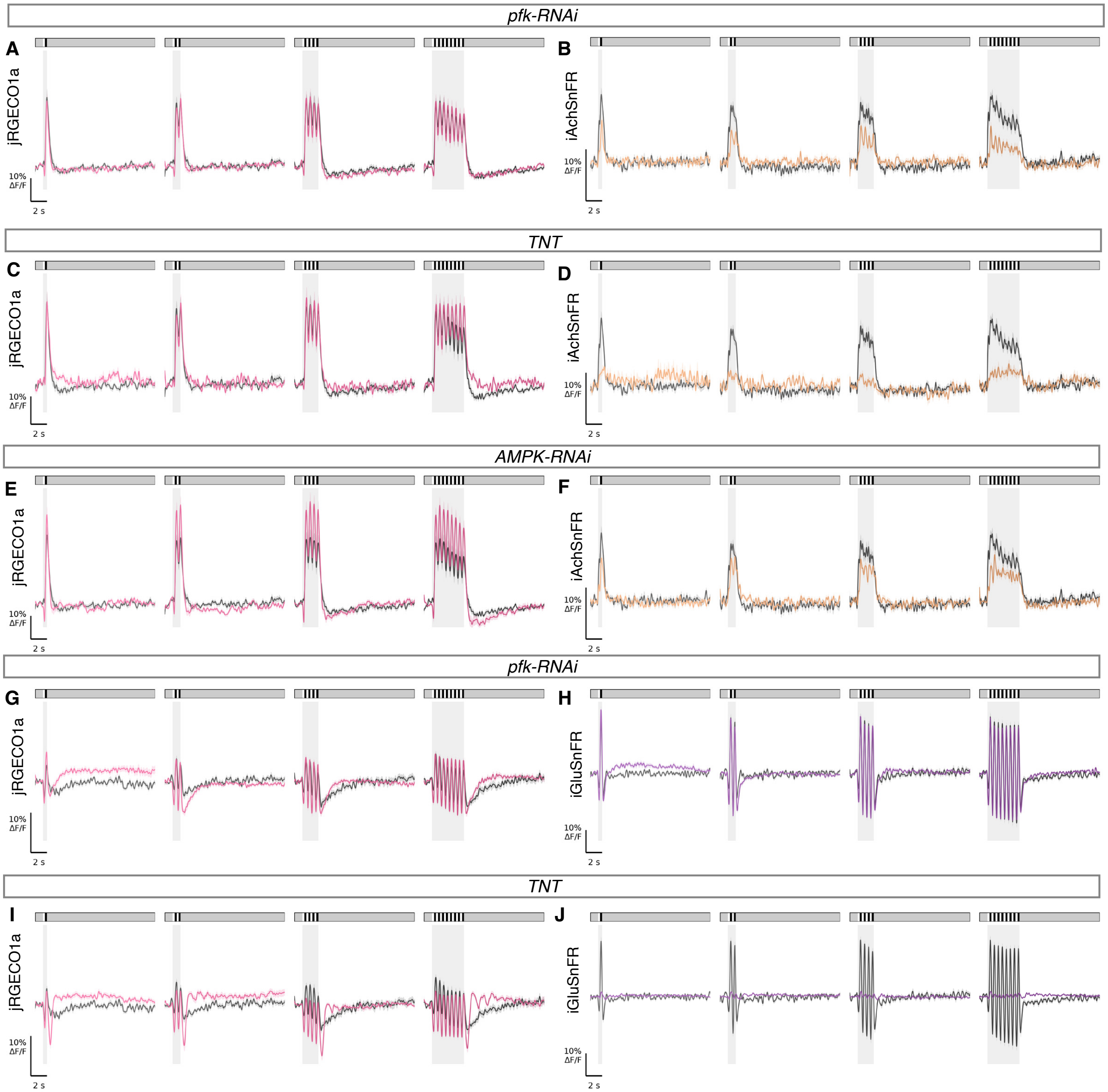


**Fig. S10. Glycolysis is selectively required for neurotransmitter release in cholinergic but not glutamatergic neurons**

(**A-F**) Mean jRGECO1a (left) and iAChSnFR (V188P) (right) responses to stimuli of four varying durations after genetic perturbations. (**A**) Mean jRGECO1a responses in control (black) and after pfk knockdown (magenta) (**B**) Mean iAChSnFR (V188P) responses in control (black) and after pfk knockdown (orange) (**C**) Mean jRGECO1a responses in control (black) and after TNT expression (magenta) (**D**) Mean iAChSnFR (V188P) responses in control (black) and after TNT expression (orange) ) (**E**) Mean jRGECO1a responses in control (black) and after AMPK knockdown (magenta) (**F**) Mean iAChSnFR (V188P) responses in control (black) and after AMPK knockdown (orange) (**G-J**) Mean jRGECO1a (left) and SF-iGluSnFR.A184V (right) responses (**G**) jRGECO1a responses in control (black) and after pfk knockdown (magenta) (**H**) Mean SF-iGluSnFR.A184V responses in control (black) and in pfk knockdown (purple) (**I**) jRGECO1a responses in control (black) and in neurons expressing TNT (magenta) (**J**) Mean SF-iGluSnFR.A184V responses in control (black) and in neurons expressing TNT (purple).
